# The Ndr/LATS kinase Cbk1 regulates a specific subset of Ace2 functions and suppresses the hyphae-to-yeast transition in *Candida albicans*

**DOI:** 10.1101/2020.07.10.198242

**Authors:** Rohan S. Wakade, Laura C. Ristow, Mark A. Stamnes, Anuj Kumar, Damian J. Krysan

## Abstract

The <u>R</u>egulation of <u>A</u>ce2 and <u>M</u>orphogenesis (RAM) pathway is an important regulatory network in the human fungal pathogen *Candida albicans*. The RAM pathway’s two most well-studied components, the NDR/Lats kinase Cbk1 and its putative substrate, the transcription factor Ace2, have a wide range of phenotypes and functions. It is not clear, however, which of these functions are specifically due to the phosphorylation of Ace2 by Cbk1. To address this question, we first compared the transcriptional profiles of *CBK1* and *ACE2* deletion mutants. This analysis indicates that, of the large number of genes whose expression is affected by deletion of *CBK1* and *ACE2*, only 5.5% of those genes are concordantly regulated. Our data also suggest that Ace2 directly or indirectly represses a large set of genes during hyphal morphogenesis. Second, we generated strains containing *ACE2* alleles with alanine mutations at the Cbk1 phosphorylation sites. Phenotypic and transcriptional analysis of these *ace2* mutants indicates that, as in *Saccharomyces cerevisiae*, Cbk1 regulation is important for daughter cell localization of Ace2 and cell separation during yeast phase growth. In contrast, Cbk1 phosphorylation of Ace2 plays a minor role in *C. albicans* yeast-to-hyphae transition. We have, however, discovered a new function for the Cbk1-Ace2 axis. Specifically, Cbk1 phosphorylation of Ace2 prevents the hyphae-to-yeast transition. To our knowledge, this is one of the first regulators of the *C. albicans* hyphae-to-yeast transition to be described. Finally, we present an integrated model for the role of Cbk1 in the regulation of hyphal morphogenesis in *C. albicans*.

**Importance:** <u>R</u>egulation of <u>A</u>ce2 and <u>M</u>orphogenesis (RAM) pathway is a key regulatory network that plays a role in many aspects of *C. albicans* pathobiology. In addition to characterizing the transcriptional effects of this pathway, we discovered that Cbk1 and Ace2, a key RAM pathway regulator-effector pair, mediate a specific set of the overall functions of the RAM pathway. We have also discovered a new function for the Cbk1-Ace2 axis; suppression of the hyphae-to-yeast transition. Very few regulators of this transition have been described and our data indicate that maintenance of hyphal morphogenesis requires suppression of yeast phase growth by Cbk1-regulated Ace2.

## Introduction

*Candida albicans* is one of the most common causes of human fungal infections, the severity of which ranges from life-threatening, invasive disease to relatively minor, but still consequential mucosal infections (1). As a component of the human mycobiome, *C. albicans* primarily colonizes the human oral cavity, gastrointestinal tract and urogenital tract (2). Accordingly, it is well-adapted to niches that differ greatly in their environmental characteristics (3). Regardless of the specific niche or site of infection, however, the ability of *C. albicans* to cause disease has been closely linked to its ability to undergo morphogenic transitions to either pseudo-hyphae or true hyphae (4). Consequently, the molecular mechanisms by which these transitions are regulated have been of keen interest to mycologists (5). Over the years, many signaling pathways have been shown to participate in this regulation including the protein kinase A pathway, MAPK signaling pathways, and the Regulation of Ace2 and Morphogenesis (RAM) pathway (6). The RAM pathway is distinguished from other, more extensively studied pathways by its apparent ability to modulate multiple transcriptional regulators (7). Here, we describe a set of experiments designed to identify the subset of Ace2 functions that are directly dependent on the RAM pathway and its kinase Cbk1.

The RAM pathway is conserved within fungi including human pathogens and its components regulate a wide variety of functions in these organisms (7). The effector components of the RAM pathway are a Ndr/LATS kinase such as Cbk1 in *Saccharomyces cerevisiae* and *C. albicans* and the zinc finger transcription factor Ace2 (7, 8). The activity of the kinase is dependent on a network of co-factors including Mob2, Kic1, Hym1, Tao3, and Sog2. The RAM pathway was initially described in the model yeast *S. cerevisiae* and its function during the cell cycle of this budding yeast have been extensively studied (8). Based on these studies, the function of Ace2 as a daughter cell specific regulator (9) of cell separation gene expression that is dependent upon phosphorylation by Cbk1 has been firmly established (10). Cbk1 phosphorylates Ace2 within the nuclear export sequence which blocks export of Ace2 from daughter cell nuclei (10). In the absence of Cbk1 phosphorylation, Ace2 localization is disrupted leading to defects in cell separation. The phosphorylation site motif for Cbk1 has been characterized biochemically and genetically leading to the following consensus sequences: HXRXXS/T and HRXXS/T (10). The histidine residue is critical for recognition and substrates are not phosphorylated in its absence. In *S. cerevisiae*, the RNA binding protein Ssd1 is the other well-characterized Cbk1 substrate (11, 12, 13) and Cbk1 phosphorylation of Ssd1 plays an important role in the regulation of translation of specific sets of genes.

In contrast, the functions of the RAM pathway and the transcription factor Ace2 in the human fungal pathogen *C. albicans* have been examined more broadly (7). In addition to their role in polarized growth and cell separation (14, 15), the RAM pathway and Ace2 affect hyphae formation (14, 15, 16), response to hypoxia (16, 17), biofilm formation (17, 18, 19), adherence to abiotic surfaces (18), susceptibility to antifungal drugs (20), cell wall structure including β-glucan masking (21, 22), and regulation of metabolic genes (16). Recently, alternative splicing has been implicated as an additional source of functional diversity for Ace2 in that one isoform appears to function as a transcription factor while the other is localized to the plasma membrane and has a role in septum dynamics (23). A genetic interaction screen reported by our group found that Cbk1 interacts with a wide range of functionally distinct genes including those noted above as well as genes required for oxidative stress tolerance and nitrogen utilization (24). In a mouse model of disseminated candidiasis, McCallum et al. found a very slight defect in survival for an *ace2*ΔΔ strain relative to WT (25). To circumvent the complicating factors of cell separation defects, our group used heterozygous mutants of *ACE2* and *CBK1* to show that both are haploinsufficient with respect to kidney burden early in the infection process (26). In summary, the RAM pathway directly or indirectly regulates a wide range of functions important to the biology and pathobiology of *C. albicans*.

One of the distinguishing features of the RAM pathway in C. albicans is that it appears to regulate multiple transcription factors. To date, three putative *C. albicans* Cbk1 substrates have been confirmed genetically or biochemically: Bcr1, Fkh2, and Ssd1. Gutierrez-Escribano et al. showed that Cbk1-mediated phosphorylation of Bcr1 is important for its full function during biofilm formation (27) and Grieg et al. demonstrated that Cbk1 phosphorylation of the Fkh2 is important for hyphal morphogenesis (28). Finally, Lee et al. reported that mutants of Ssd1 lacking Cbk1 phosphosites failed to degrade *NRG1* mRNA during hyphal morphogenesis, providing a mechanism for the profound inability of *cbk1*ΔΔ mutants to generate hyphae (29).

In contrast to *S. cerevisiae*, Ace2 has not been experimentally confirmed to be a Cbk1 substrate in *C. albicans*. A large-scale phosphoproteomic analysis of *C. albicans* during hyphae formation showed that two of the three putative Cbk1 phosphorylation sites contain phosphates under those conditions (30). Although there is little doubt that Ace2 is regulated by Cbk1, we were interested in identifying Cbk1-dependent functions of Ace2. Here, we describe the generation of strains with the Cbk1 phosphorylation sites mutated to non-phosphorylatable alanine residues (*ace2-2A* and *ace2-3A*). As described below, our observations indicate that, although both Cbk1 and Ace2 have a wide range of functions important to the biology of *C. albicans*, a very specific set of these functions are directly regulated by the phosphorylation of Ace2 by Cbk1, including maintaining a balance between hyphal and yeast transcriptional programs (31).

## Results

### Cbk1 regulates a relatively small proportion of Ace2-dependent genes during yeast phase growth

The RAM pathway and Ace2 have been the subject of previous transcriptional profiling experiments using microarray technology. Mulhern et al. profiled *ace2*ΔΔ mutants during both yeast phase growth and serum-induced hyphal growth (16). In addition, Song et al. examined the effect of deleting *MOB2*, the Cbk1 binding partner and activator, under similar conditions (20). Since RNA sequencing had not been used in either instance, we decided to compare the transcriptional profiles of *ace2*ΔΔ and *cbk1*ΔΔ using this technique. We were particularly interested in identifying the set of genes that is regulated by both Cbk1 and Ace2.

During yeast phase growth at 30°C in rich, yeast peptone dextrose (YPD), the expression of 526 genes is altered by at least 2-fold (adjusted *P* <0.05, full set of genes listed in Table S3) in the *ace2*ΔΔ mutant (Fig. 1A). The total number of genes affected by deletion of *ACE2* observed in our data set is very similar to that reported by Mulhern et al. (reference 16, 363 with FDR 0.23%; 645 with FDR 0.9%). In our experiment, more genes (475) were downregulated than were upregulated (51) in the *ace2*ΔΔ mutant; again, this was similar to Mulhern et al. (16) as were the GO terms and gene groups affected.

**Figure 1.**
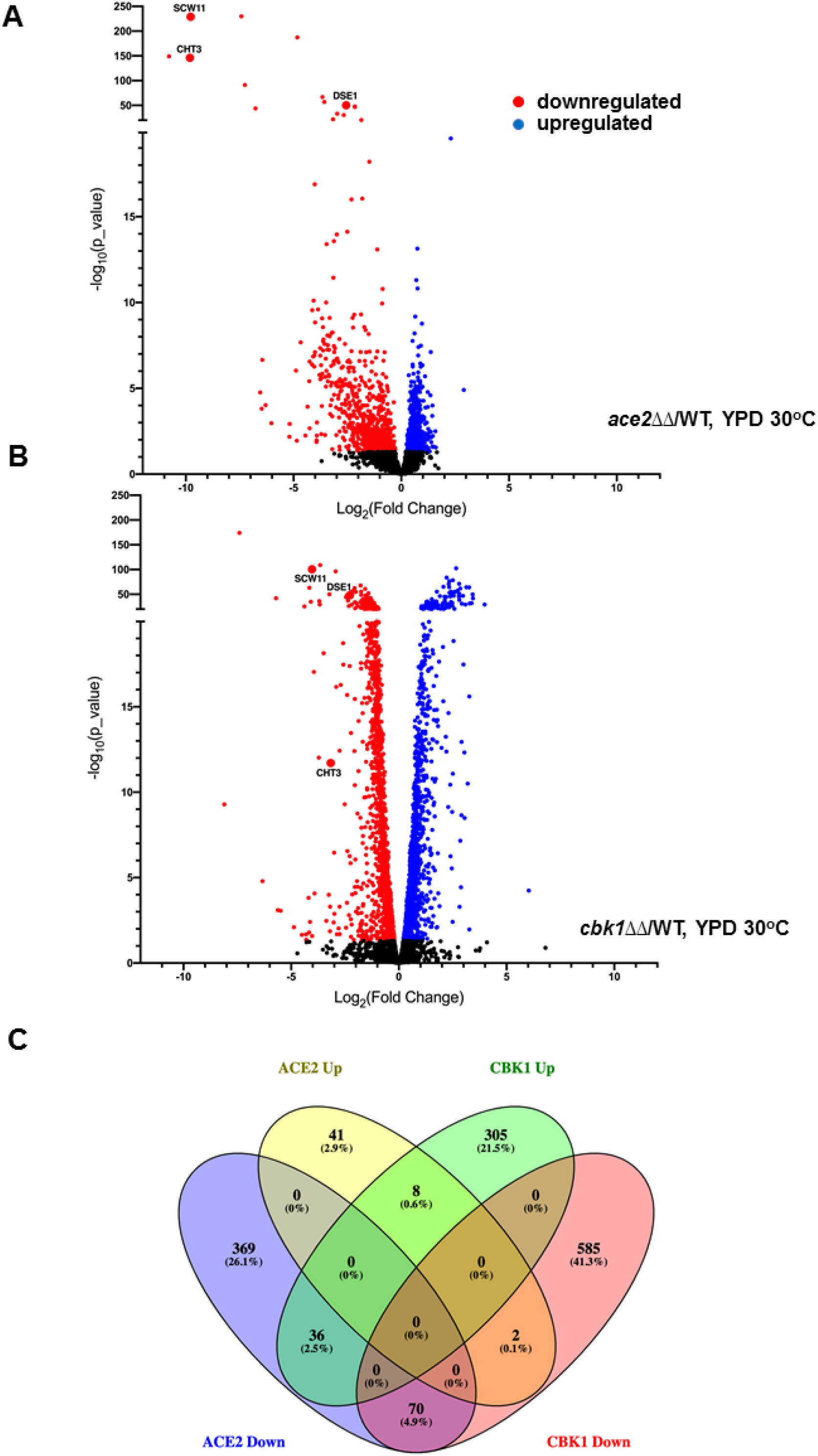
Ace2 and Cbk1 affect the expression of large sets of genes during yeast phase growth but only a fraction of the genes is regulated in common. Volcano plots of the gene expression profiles of *ace2*ΔΔ (A) and *cbk1*ΔΔ (B) relative to wild type under exponential growth in YPD at 30°C with genes showing statistically significant (adjusted *P* < 0.05) changes in expression. (C) Venn diagram of overlap between genes downregulated by log_2_ 1 in *ace2*ΔΔ and *cbk1*ΔΔ mutants.

We also isolated RNA from *cbk1*ΔΔ cells grown under the same conditions. Somewhat surprisingly, 1006 genes were differentially expressed in *cbk1*ΔΔ relative to WT during logarithmic yeast phase growth with 657 genes downregulated by at least 2-fold and 349 genes upregulated to the same extent (Fig. 1B, Table S4). These data indicate that Cbk1 has a significant effect on gene expression in *C. albicans*. As shown in the Venn diagram in Fig. 1C, only 70 genes are downregulated in both *ace2*ΔΔ and *cbk1*ΔΔ mutants (9.5% of Cbk1 and 13% of Ace2 sets). A list of the specific genes in each category of the Venn diagrams is provided in Supplementary Table 6. The overlap between the genes upregulated in *cbk1*ΔΔ and *ace2*ΔΔ strains is even less (8 genes). Thus, Cbk1 and Ace2 directly or indirectly affect the expression of significant number of genes during yeast phase growth, but only a small fraction of these genes appears to dependent on the function of both proteins.

Consistent with previous single gene analyses of both Cbk1 and Ace2, cell septum degrading enzymes *CHT3* and *SCW11* (Fig. 1A&B) are among the set of genes downregulated in both mutants. Of the remaining genes, ribosome/RNA processing (FDR 0.00%) was the only GO term that was enriched in the set of genes differentially regulated in both *cbk1*ΔΔ and *ace2*ΔΔ. These data, therefore, suggest that Cbk1-Ace2 primarily functions to regulate septum degradation during yeast growth while both Ace2 and Cbk1 have significant, wide-ranging effects on gene transcription that are independent of one another.

### Ace2 directly or indirectly represses the expression of a large set of genes during hyphal morphogenesis in Spider medium

We also characterized the effect of an *ace2*ΔΔ mutation on gene expression during hyphae morphogenesis; *cbk1*ΔΔ mutants are unable to form hyphae (14, 19, 20) and, thus, we did not perform expression analysis under hyphae-inducing conditions. Strains lacking *ACE2* form hyphae in liquid culture in response to a variety of standard inducing conditions and agents. Of these, Spider medium (SM) at 37°C is the only inducing condition under which *ace2*ΔΔ mutants differ from WT (32); *ace2*ΔΔ mutants form hyphae modestly slower than WT. We, therefore, used this inducing condition rather than serum-induction that was employed by Mulhern et al (16). In addition, we harvested cells after 4 hours of induction, a time point where over 75% of the *ace2*ΔΔ have formed hyphae and Ace2 protein levels have peaked by western blot analysis (32).

In hyphae-inducing conditions (SM, 37°C), a total of 575 genes were differentially expressed by at least 2-fold in the *ace2*ΔΔ mutant relative to the reference strain (Fig. 2A, Table S5). Interestingly, the set of genes was almost equally split between those with increased expression (283) and those with decreased expression (292). This is in distinct contrast to the relatively modest number of genes upregulated in *ace2*ΔΔ mutants during yeast phase growth (Fig. 1A&C). This is also quite different from the number of up-regulated genes (18 genes, 2-fold decreased) observed by Mulhern et al. (16) during hyphal induction with serum-containing medium. Nearly two-thirds of the genes upregulated in the *ace2*ΔΔ are involved in metabolism of carbon or nitrogen while, as with during yeast phase growth, approximately 40% of the genes are related to RNA/ribosome function.

**Figure 2.**
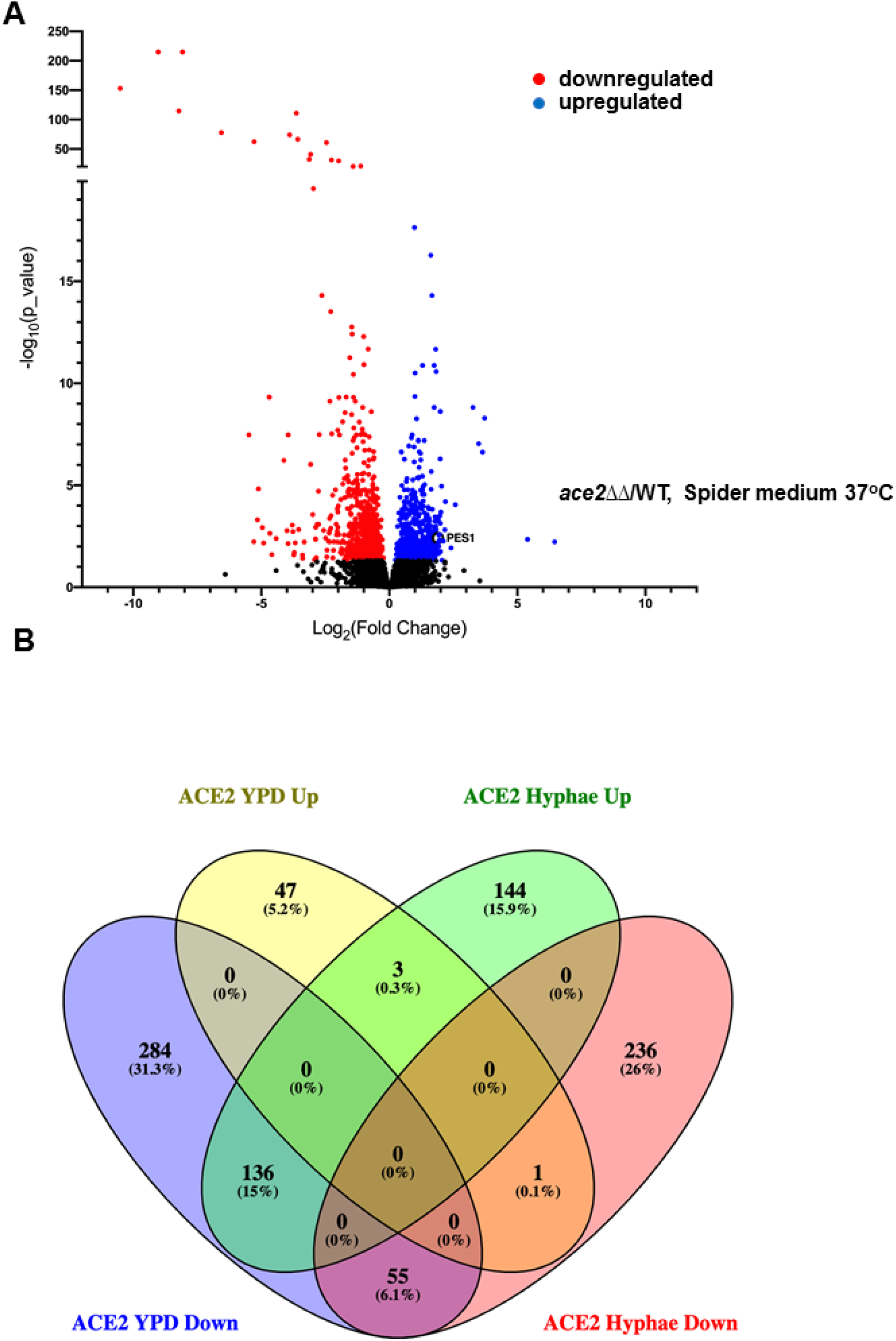
Ace2 directly or indirectly represses a large set of genes during hyphal morphogenesis. (A) Volcano plot of the gene expression profiles of *ace2*ΔΔ cells relative to wild after 4 hours in Spider medium (SM) at 37°C with genes showing statistically significant (adjusted *P* < 0.05) changes in expression of ± log_2_ 1. (B) Venn diagram of differentially expressed genes (adjusted *P* <0.05) for an *ace2*ΔΔ relative to wild type under yeast phase and hyphal growth.

It is important to note that very few genes are concordantly regulated by Ace2 during yeast and hyphal growth. Specifically, 54 genes are downregulated in both YPD and SM while while only 3 genes are upregulated in both conditions. Although the canonical Ace2 targets *SCW11*, *CST1*, and *DSE3* are among the sets of genes downregulated in both conditions, the only GO terms that are significantly enriched among the concordantly regulated genes are carbon utilization (*ICL1*, *MLS1*, *PCK1*, *PDK2*, *PYC2*, and *SFC1*; FDR 0.00%) and acetate metabolism (*CTN1*, *ICL1*, *MLS1*, *SFC1*, FDR 0.00%). Indeed, 136 genes are downregulated in *ace2*ΔΔ mutants in YPD at 30°C but are upregulated during hyphal induction in SM at 37°C. Once again, this set of genes is dominated by RNA/ribosome processing and metabolic functions. The large number of discordantly regulated genes emphasizes the context dependent effect that Ace2 appears to have on *C. albicans* gene expression. Overall, these data indicate that Ace2 directly or indirectly affects the expression of many genes during hyphal morphogenesis in SM and that it appears to play a previously unappreciated role in repressing the expression of many genes under hyphae-inducing conditions.

### Mutation of the Cbk1 phosphorylation sites in Ace2 by CRISPR/Cas9-mediated genome editing

Previous phenotypic characterization of *ACE2* has shown that the *C. albicans* and *S. cerevisiae* orthologs have similar functions during yeast phase growth and, in particular, are important for proper cell separation. As depicted in Fig. 3A, *S. cerevisiae* Ace2 has four Cbk1 phosphorylation motifs with two in the nuclear export sequence (NES), one N-terminal to that site, and one near the DNA binding domain in the C-terminal region of the protein (10). The cell cycle functions of *Sc*Ace2 are dependent on S122, S137 and S436, indicating that *Sc*Cbk1 is not only required for concentration of *Sc*Ace2 in the nucleus through S122/S137 phosphorylation but also must contribute additional activation functions by phosphorylating S436 (10).

**Figure 3.**
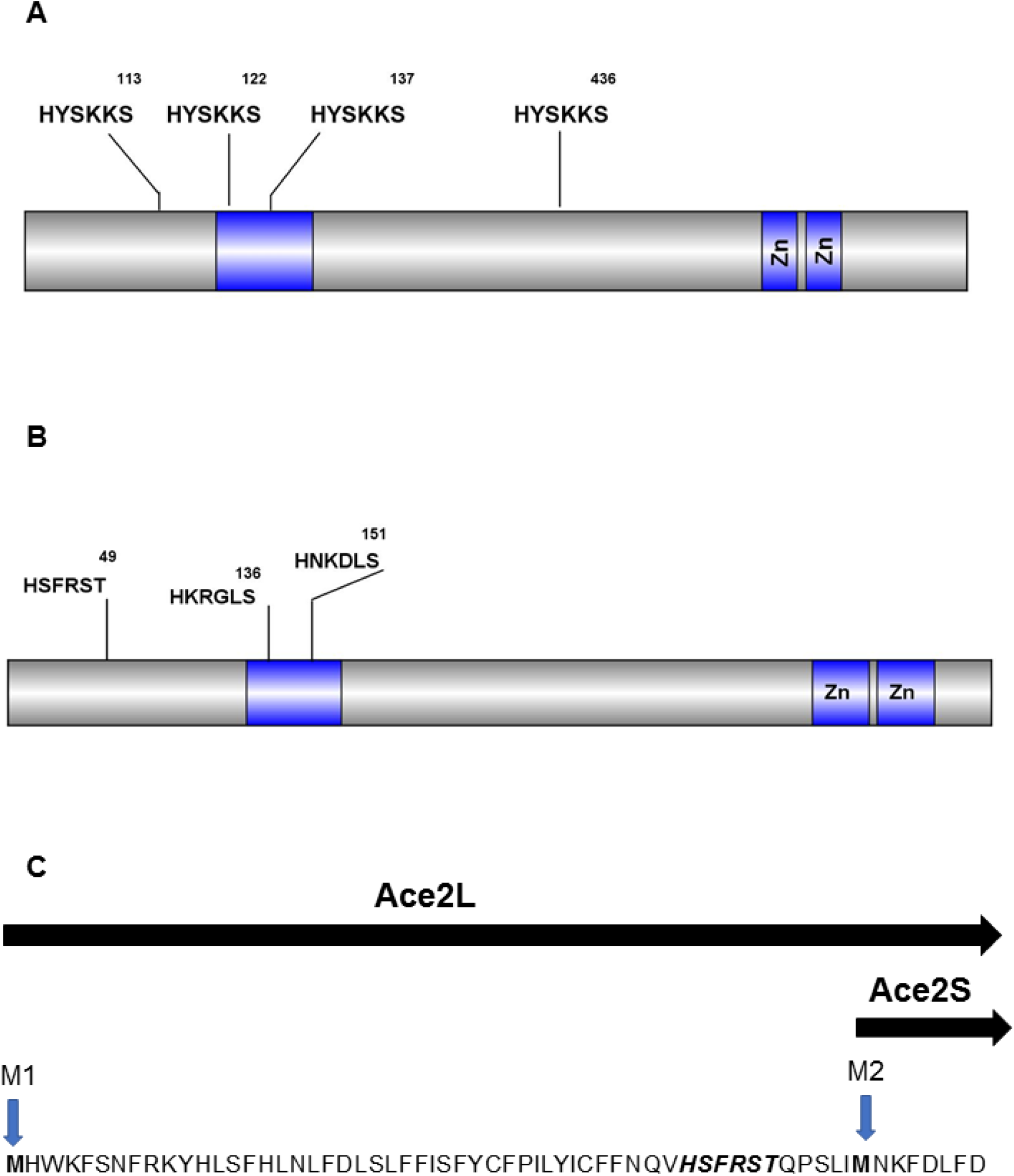
Schematic of putative Cbk1 phosphorylation sites and alternative isoforms of CaAce2. Schematic representation of Cbk1 phosphorylation motifs in *S. cerevisiae* (A) and *C. albicans* (B) Ace2. (C) Sequence of the N-terminus of CaAce2 showing the two alternative AUG sites for initiation of translation and the resulting long (Ace2L) and short (Ace2S) as formulated by Calderón-Noreña et al. (23).

Ace2 (we will use Ace2 to refer to the *C. albicans* ortholog and *Sc*Ace2 to refer to the *S. cerevisiae* ortholog for the remainder of the text) lacks the C-terminal Cbk1 phosphorylation motif but has sites in the NES and in the N-terminal region (Fig. 3B). Willger et al. performed a comprehensive phosphoproteomic study of *C. albicans* during hyphae induction and confirmed that Ace2 S136 and S151 are phosphorylated (30). There is evidence that, in the SC5312 strain background (23), Ace2 has two isoforms depending on the translational start site (Fig. 3B). The N-terminal Cbk1 phosphosite is between the two start sites and, thus, would only be expected to be present in the Ace2L form that is proposed to associate with the plasma membrane and play a role in septin dynamics. Ace2S is the isoform that localizes to the nucleus and regulates the expression of cell separation genes (23).

To assess the role of Cbk1 phosphorylation in Ace2 function, we used a CRISPR/Cas9 approach to mutate the serine and threonine Cbk1 consensus phosphorylation sites of the endogenous *ACE2* to alanine (33). The details of this strain construction are described in the materials and methods. In this way, we constructed a strain in which the only *ACE2* allele lacked the Cbk1 phosphorylation sites in the NES (S136A, S151A, *ace2*-2A) and a strain that lacks all three of the Ace2, Cbk1 phosphorylation sites (S49A, S136A, S151A, *ace2*-3A). The initially isolated phosphosite mutants were heterozygous at the *ACE2* locus with a wild type allele and the desired mutant allele. We deleted the wild type allele using standard homologous recombination and confirmed that the only remaining allele contained the S/T-A mutations by Sanger sequencing. Thus, the resulting strains are heterozygous at the *ACE2* locus and, if the Cbk1 phosphorylation sites are required for function, the strains would be expected to have phenotypes that are similar to an *ace2*ΔΔ homozygous deletion. In our phenotypic characterization data presented below, *ace2*ΔΔ is used as the control; as shown in Supplementary Data, a*ce2*Δ/*ACE2* strains show no haploinsufficiency and thus phenotypic changes are due to the mutations and not to changes in gene copy number (Fig. S1).

### Phospho-acceptor amino-acids at Ace2 consensus Cbk1 substrate motifs are required for normal cell separation in *C. albicans* during yeast phase growth

A phenotype of *ACE2* and *CBK1* mutants that is conserved across all yeast species studied to date is decreased cell separation due to reduced septum degradation (8, 9, 15). Therefore, we compared the cell separation characteristics of the *ace2*-2A and *ace2*-3A mutants to wild type and *ace2*ΔΔ mutant strain under standard yeast culture conditions (30°C, YPD media). The harvested cells were fixed and examined by light microscopy to determine the relative proportion of small, medium, and large cell clusters (Fig. 4A). Consistent with previous literature (8, 9, 15), WT cultures were dominated by small clusters (1-2 cell) while the *ace2*ΔΔ mutant cultures were almost entirely comprised of large cell clusters (>5 cells, Fig. 4A). The *ace2*-2A mutant showed an intermediate phenotype with increased numbers of medium and large cell clusters that were not changed significantly by mutation of the non-NES Cbk1 site (*ace2*-*3A*). As previously found for *ace2*ΔΔ mutants (8, 9, 15), ultrasonication of the cell clusters formed by the *ace2*-*2A* and *ace2*-*3A* mutants did not alter the size of the clusters which is consistent with a failure of the cells to separate and is inconsistent with non-covalent aggregation (data not shown).

**Figure 4.**
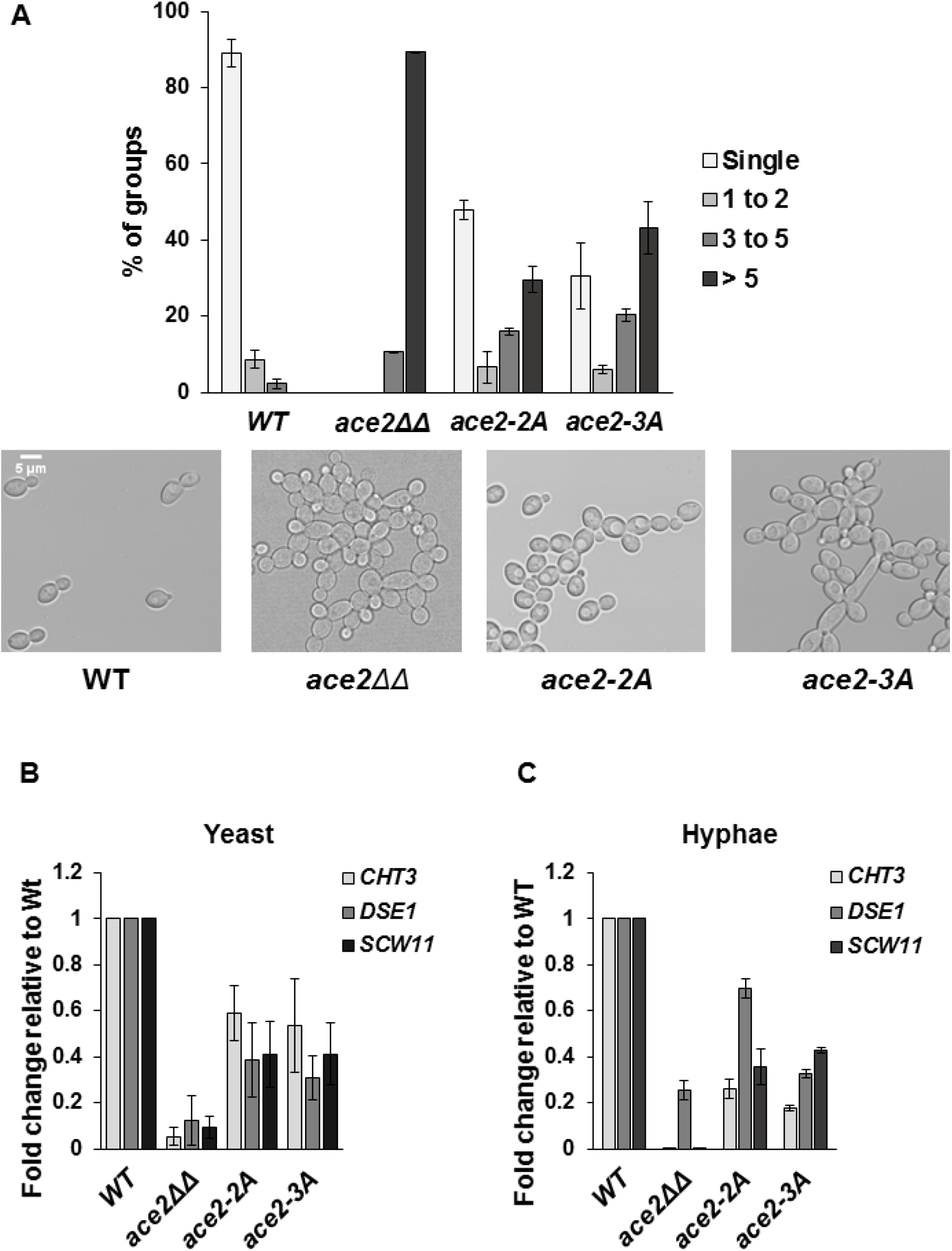
Cbk1 phospho-acceptor sites for cell separation and expression of septum-degrading enzymes. (A) Exponential phase cultures of the indicated strains in YPD at 30°C were harvested and the distribution of cell aggregates was determined by bright field microscopy with the cells binned into the indicated groups. Representative photomicrographs of fields are shown for each strain. The bars indicate mean of three independent experiments (N ≥ 150) with SD indicated by error bars. The differences in cell aggregates was statistically different from wild type for all mutant strains (Student’s t test, *P*<0.05) (B) The expression of the indicated Ace2 genes was determined by quantitative RT-PCR. The expression was normalized to *ACT1* by the ΔΔ^ct^ method and normalized to WT. The bars indicate mean values for each strain with error bars indicating the SD of three independent experiments with technical replicates. The expression of each gene in the indicated mutants was statistically different than WT (Student’s t test, *P*<0.05).

As noted above, this phenotype is attributed to the decreased expression of septum-degrading enzymes such as *CHT3*, *DSE1*, and *SCW11* in the *ace2*ΔΔ mutant (Fig. 4B&C). The expression of these genes is also decreased in both *ace2*-*2A* and *ace2*-*3A* mutants during both yeast and hyphal cell growth (Fig. 4B&C). As with the cell separation data, mutations within the NES site are responsible for the majority of the observed effects on septum-degrading gene expression. This is consistent with the findings that Ace2S form of the protein is responsible for transcriptional regulation of cell separation genes while the site in the Ace2L form does not play a significant role in the expression of these genes (23). These data are consistent with previous studies in *S. cerevisiae* indicating that Cbk1 phosphorylation of the NES in Ace2 is required for expression of cell septum-degrading enzymes and proper cell separation (10).

### Cbk1 phospho-acceptor sites in a putative nuclear export signal site are required to concentrate Ace2 to daughter cell nuclei in *C. albicans*

In *S. cerevisiae* daughter cells, phosphorylation of *Sc*Cbk1 consensus motifs in the NES of *Sc*Ace2 prevents its export and, thereby, concentrates *Sc*Ace2 within the nuclei of new daughter cells while it is relatively excluded from mother cell nuclei (10). Mutations that prevent *Sc*Cbk1-mediated phosphorylation of *Sc*Ace2 result in the transcription factor localizing to both daughter and mother cell nuclei (10). To determine if a similar process occurs in *C. albicans*, we tagged the C-terminus of *ACE2* in wild type and *ace2*-*2A* strains with green fluorescent protein (GFP); we and others have shown previously that WT alleles retain function (15, 34) and we confirmed that the phenotypes of the *ace2*-*2A*-GFP strains did not differ from the parental strain (Fig. S2).

Consistent with previous reports, WT Ace2 localized to the daughter cell in budding yeast phase (Fig. 5A). In contrast, *ace2*-*2A*-GFP signal was present in both cells of mother-daughter pairs (Fig. 5B). This same pattern of mislocalization is consistent with that reported by the Weiss lab for strains containing *ScACE2* alleles lacking Cbk1 phosphoacceptor sites in the NES domain of the protein (10). This localization pattern also provides an explanation for the intermediate cell separation phenotype (relative to the *ace2*ΔΔ strain) observed for the *ace2*-*2A* strains because some cells localized Ace2 to the daughter cell nuclei whereas in the null mutant there is a complete absence of protein. We attempted to characterize the effect of the *ace2-2A* allele on localization in hyphae but the relatively few wild type hyphae that show the canonical Ace2 localization to the distal most nuclei of the hyphae prevented our ability to make firm conclusions (Wakade and Krysan, unpublished results). These data indicate that, as in *S. cerevisiae*, Cbk1 phosphorylation of Ace2 within the putative NES is required to concentrate Ace2 in daughter cell nuclei during yeast phase growth.

**Figure 5.**
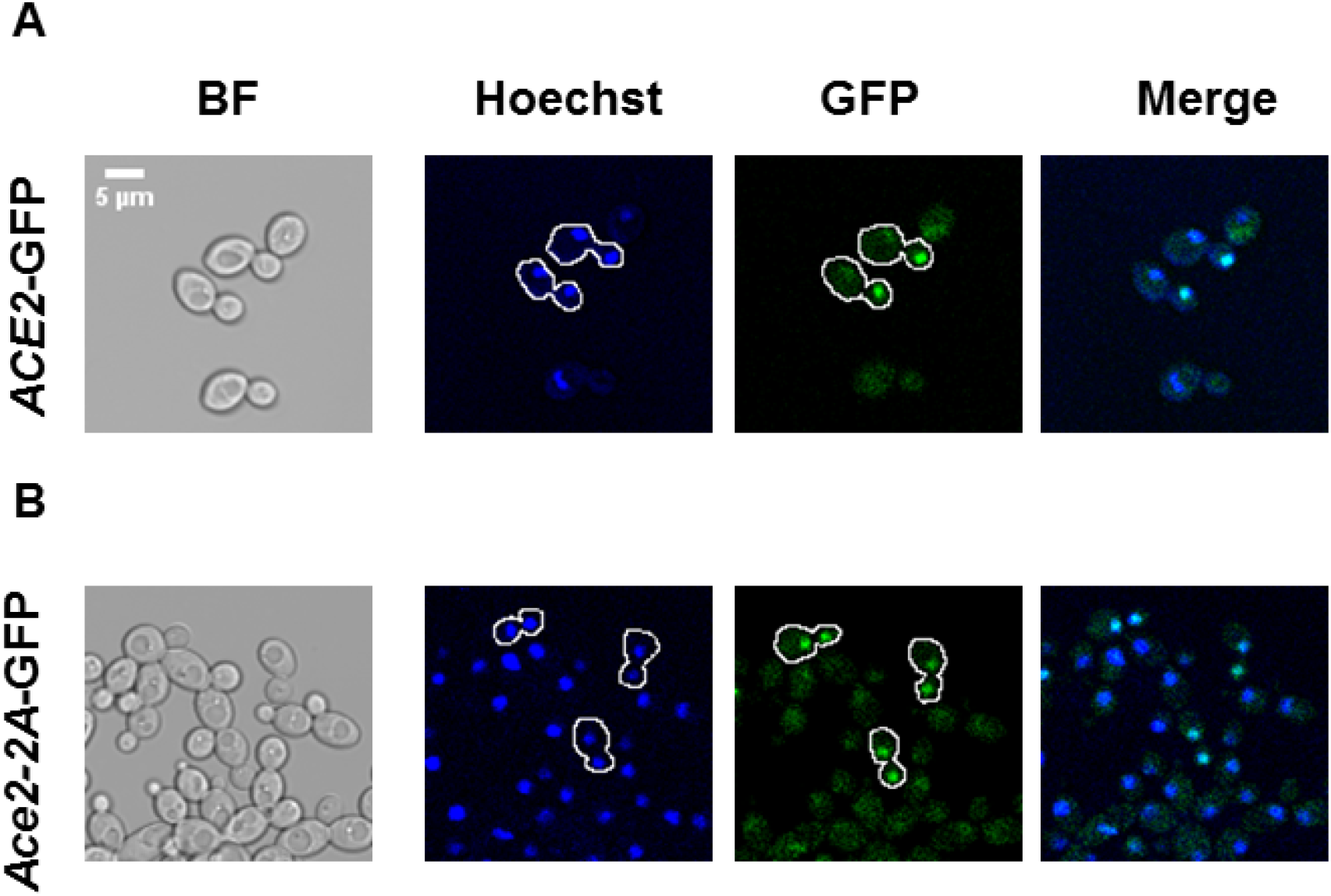
Cbk1 phospho-acceptor sites in the putative nuclear export site of Ace2 are required for exclusive localization to daughter cell nuclei in yeast phase cells. Exponential cells for the *ACE2*-GFP (A) and *ace2-2A-*GFP were harvested and stained with Hoechst before imaging with the indicated channels. The outlines of the indicated cells were generated to highlight mother-daughter pairs. Scale bar 5 μm.

### Cbk1 phosphorylation of Ace2 affects susceptibility to chitin-targeted cell wall stressors

Ace2 function has been implicated in maintaining cell wall homeostasis through multiple pathways (7). First, *ace2*ΔΔ mutants are resistant to the chitin binding molecule, calcofluor white (CFW). This is most likely due to an increase in the chitin content of the wall in the region of septum due to decreased expression of chitinases. Consistent with our observation that the *ace2*-*2A/3A* mutants have decreased expression of *CHT3*, they are also resistant to concentrations of CFW that inhibit the growth of WT cells (Fig. 6A); similarly, both *ace2*ΔΔ and the *ace2*-*2A*/*3A* mutants are resistant to Congo Red, a molecule that interacts with both chitin and glucan components of the cell wall (Fig. 6B). Ace2 has also been shown to affect the mannoprotein layer of the cell wall through a separate signaling pathway involving Cek1 (21); a manifestation of this function is the increased susceptibility of the *ace2*ΔΔ mutant to the glycosyl-transfer inhibitor, tunicamycin (Fig. 6C). The *ace2*-*2A/3A* mutants grow similar to wild type cells at concentrations of tunicamycin that inhibits *ace2*ΔΔ growth, indicating that this function of Ace2 is independent of Cbk1. Thus, the cell wall functions regulated by Cbk1-Ace2 appear to be mainly limited to cell septum-related processes while other cell wall functions of Ace2 are independent of Cbk1.

**Figure 6.**
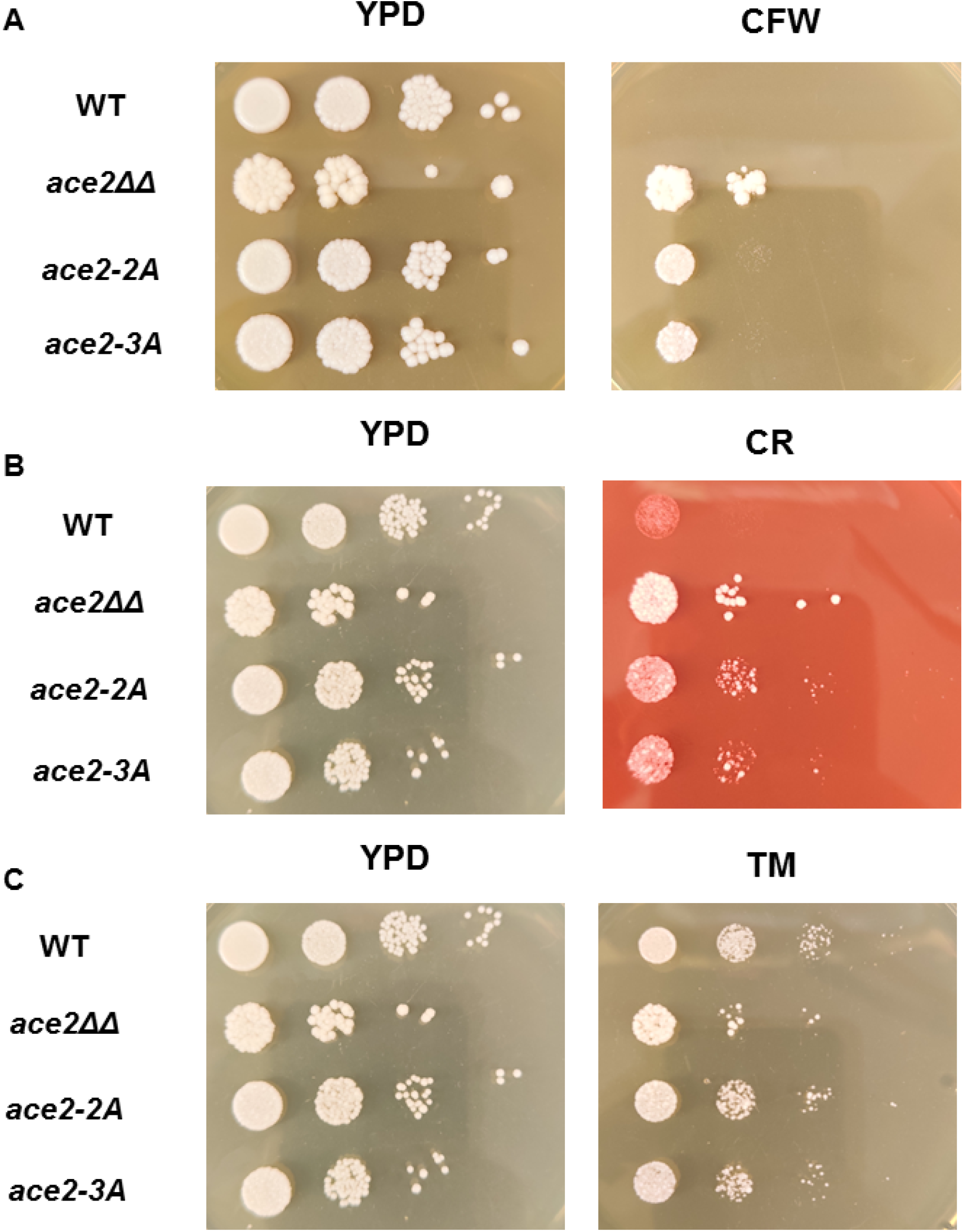
Cbk1 phospho-acceptor site mutants of *ACE2* are resistant to chitin-binding molecules but not tunicamycin. A ten-fold dilution series (0.1 OD_600_ initial cell density) was plated on YPD as well as YPD containing calcofluor white (50 μg/mL, A), Congo red (400 μg/mL) and tunicamycin (4 μg/mL, C). The plates were incubated for 3 days at 30°C before being photographed. The images are representative of two to three independent replicates.

### Cbk1 phosphoacceptor mutants have context-specific effects on the role of Ace2 under filament-inducing conditions

The role of Ace2 during *C. albicans* hyphal morphogensis is dependent on the environmental conditions that induce the morphological transition. Under non-inducing conditions, cultures of *ace2*ΔΔ mutants have increased numbers of pseudohyphal cells (15, 16). As a result, the colony morphology of *ace2*ΔΔ mutants on YPD plates at 30°C takes on a wrinkled or scalloped appearance while wild type strains form typical smooth colonies (Fig. 7A). This well-described phenotype suggests that Ace2 may play a role in suppressing the hyphal morphogenesis program in daughter yeast cells during non-inducing conditions. As shown in Fig. 7A, the colony morphology of *ace2*-2A/*ace2*-3A mutant is similar to the smooth surface of WT strains, indicating that Cbk1 plays a minor role in this function of Ace2.

**Figure 7.**
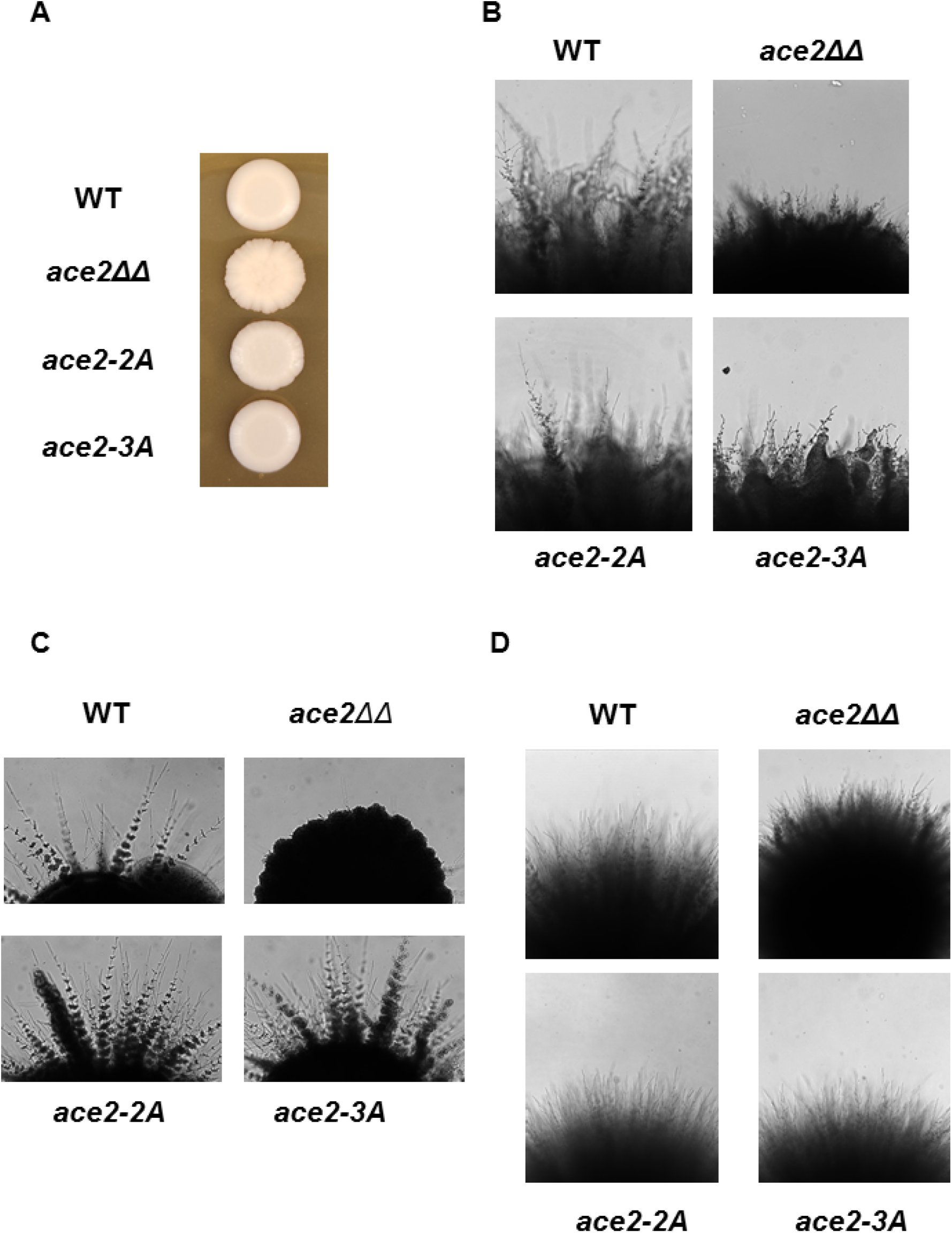
Cbk1-phosphorylation of Ace2 has a modest effect on hyphae formation under hypoxic conditions. (A) The scalloped colony morphology of the *ace2*ΔΔ mutant is not phenocopied by the Cbk1 phospho-acceptor site mutants *ace2-2A* or *ace2-3A* after 3 days at 30°C on YPD. (B) Cbk1 phospho-acceptor site mutants *ace2-2A* or *ace2-3A* have modestly reduced hyphae formation under hypoxic conditions (0.1% O_2_; surface inoculation, YPS; 2 days). The Cbk1 phosphorylation sites of Ace2 are not required for hyphal morphogensis under embedded conditions (YPS) at 25°C (C) or 37°C (D) but *ACE2* is required for hyphae formation at 25°C.

Both the Butler and Ernst groups have shown that Ace2 is required for hyphae formation under hypoxic conditions (16, 17). As shown in Fig. 7B, in the SN strain background, Ace2 is not absolutely required for hyphae formation under hypoxic conditions but the *ace2*ΔΔ mutant strain clearly is deficient in hyphae formation relative to the reference strain. The *ace2*-2A and *ace2*-3A mutants also undergo less robust hyphal morphogenesis relative to the reference strain under hypoxic conditions, suggesting that Cbk1-Ace2 pair play a modest role during hypoxia-induced hyphae formation. Mulhern et al. (16) also reported that *ace2*ΔΔ mutants are dramatically deficient for hyphae formation when embedded in YPS at 25°C and we observed the same phenotype for our *ace2*ΔΔ strains (Fig. 7C). Interestingly, the *ace2*-*2A* and *ace2*-*3A* mutants formed hyphae robustly under these conditions (Fig. 7C). Consistent with the data of Mulhern et al. (16), WT as well as the *ace2*-*2A*/*ace2*-*3A* mutants formed hyphae when embedded in YPS and incubated at 37°C instead of 25°C (Fig. 7D). These data indicate that the role of Ace2 in supporting the formation of hyphae is largely independent of Cbk1.

### Identification of a novel Ace2 function during hyphal morphogenesis: Cbk1-dependent suppression of the hyphae-to-yeast transition

We had previously reported that *ace2*ΔΔ mutants undergo hyphal morphogenesis in liquid media containing a variety of inducing agents (32), although the tempo of hyphae formation is modestly slower in SM. Upon re-examination of this process, we noted that *ace2*ΔΔ strains developed lateral yeast cells at sub-apical segments of the hyphal filament before they were evident on WT hyphae (Fig. 8A). In addition, we observed yeast phase budding from the mother cells from which the hyphal structure emerged, indicating that, in the absence of *ACE2*, hyphal-mother cells had re-entered the cell cycle earlier than WT cells (Fig 8A). As hyphae mature, a hyphae-to-yeast transition eventually occurs leading to the emergence of yeast buds at sub-apical cell compartments within the hyphae (31, 35). Quantification of this observation confirmed that at 4 hours induction, only 10% of WT hyphae displayed lateral yeast cells at sub-apical segments whereas 35% of *ace2*ΔΔ mutants had formed lateral yeast cells (Fig. 8B). Finkel et al. had previously reported that Snf5 regulates *ACE2* expression in SM and our inspection of photomicrographs of *snf5*ΔΔ mutants suggested that there may be increased lateral yeast formation in these strains as well (18). Consistent with that assessment, 80% of hyphae formed by the *snf5*ΔΔ mutant have lateral yeast at a time point when wild type cells have only 10% (Fig. 8B). Our data indicate that Ace2 and Snf5 repress lateral yeast formation and further suggest that Snf5 and Ace2 are required to ensure that the transition from hyphae to yeast does not occur prematurely. In addition, it appears that *ace2*ΔΔ is required to prevent hyphal mother cells from re-entering the cell cycle prematurely.

**Figure 8.**
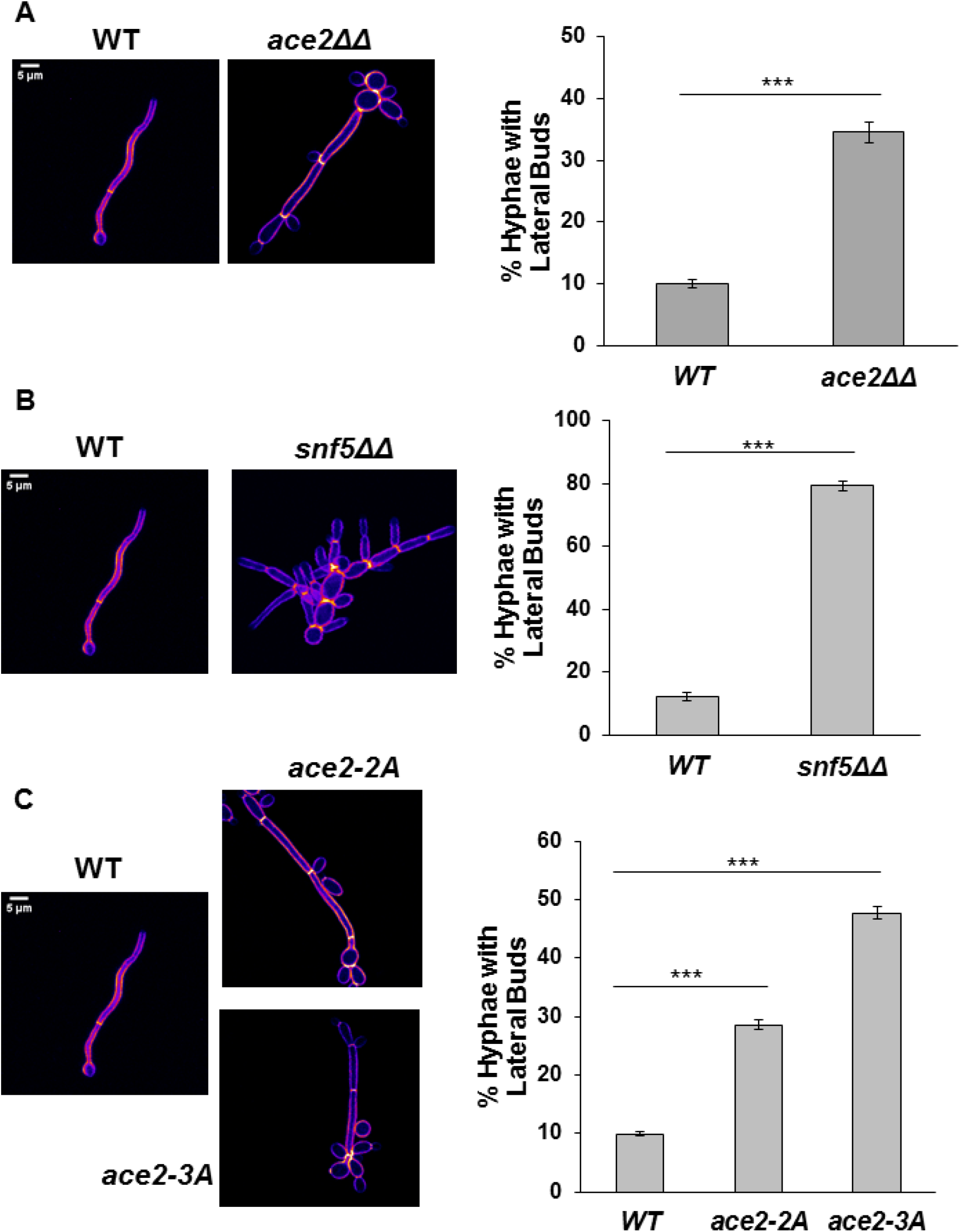
Ace2 and Snf5 suppress the hyphae-to-yeast transition in a Cbk1-dependent manner. The indicated strains were induced to form hyphae in Spider medium (SM) at 37°C for 4 hours. The cells were fixed and the percentage of hyphae with yeast buds at subapical compartments was determined. The bars indicate the mean of three independent experiments (N ≥ 100) with SD indicated by the error bars. Representative images of calcofluor white stained cells of WT and *ace2*ΔΔ (A), *snf5*ΔΔ (B), *ace2-2A/3A* (C) are shown. Bar 5 μm.

We next asked whether Cbk1 was required for the ability of Ace2 to suppress lateral yeast cell formation during the maintenance phase of hyphae formation. As shown in Fig. 8C, the *ace2*-*2A* and *ace2*-*3A* cells showed increased lateral bud cell formation indicating that this Ace2 function is likely to be Cbk1-dependent. The extent of lateral yeast cell formation is also increased in the *ace2*-*3A* mutant relative to the *ace2*-*2A* mutant (Fig. 8C). Phosphorylation at S49 is only relevant if the Ace2L isoform plays a role (Fig. 3B); if Ace2S was the operative isoform then the phenotypes of *ace2*-*2A* and *ace2*-*3A* should be identical (23). Taken together, these data indicate that the Cbk1-Ace2 axis functions to suppress the hyphae-yeast transition during the maintenance phase of hyphal morphogenesis in *C. albicans*.

### Expression of *PES1*, a regulator of the hyphae-to-yeast transition, is increased in *ace2*ΔΔ and *ace2* phosphosite mutants

The hyphae-to-yeast transition has not been studied to nearly the same extent as the yeast-to-hyphae transition (31,35). In pioneering work, the Koehler lab found that *PES1*, a pescadillo homolog, is required for the hyphae-to-yeast transition and is essential in yeast phase cells but not during hyphal phase growth (31). Conversely, increased expression of *PES1* induces increased lateral yeast formation. We, therefore, examined our RNA sequencing data set to determine the expression level of *PES1* in *ace2*ΔΔ. Indeed, *PES1* expression is increased 3.8-fold in hyphal *ace2*ΔΔ relative to wild type but is actually reduced in yeast *ace2*ΔΔ cells (Tables S3 & S5); the same trend was observed by Mulhern et al. in their transcriptional profile of *ace2*ΔΔ mutants (16). To confirm this observation, the expression of *PES1* was measured by qRT-PCR in WT and *ace2*ΔΔ strains after 3 hours of induction with SM (Fig. 9A). The expression of *PES1* was increased in the *ace2*ΔΔ strain relative to wild type, confirming the genome-wide expression profiling data and strains containing Ace2 Cbk1-phosphosite mutations also have increased expression of *PES1* (Fig. 9A). Consistent with the increased lateral yeast formation in the *ace2-3A* strain relative to *ace2-2A* mutants, the *ace2-3A* mutant shows a trend toward increased *PES1* expression relative to the *ace2-2A* mutant (Fig.8B). Interestingly, the archetypal hyphal specific gene *HWP1* is expressed at increased levels in the *ace2*ΔΔ deletion and the Cbk1 phosphosite mutants as well (Fig. 9C), indicating that, despite increased development of lateral yeast, the hyphal transcriptional program is still strongly induced. Thus, the expression of well-characterized reporter genes for both the yeast-to-hyphae and the hyphae-to-yeast transcriptional programs is upregulated in *ace2* mutants. This expression profile suggests that the Cbk1-Ace2 axis plays an important role in regulating the balance between morphogenic transcriptional programs during hyphae formation in *C. albicans*.

**Figure 9.**
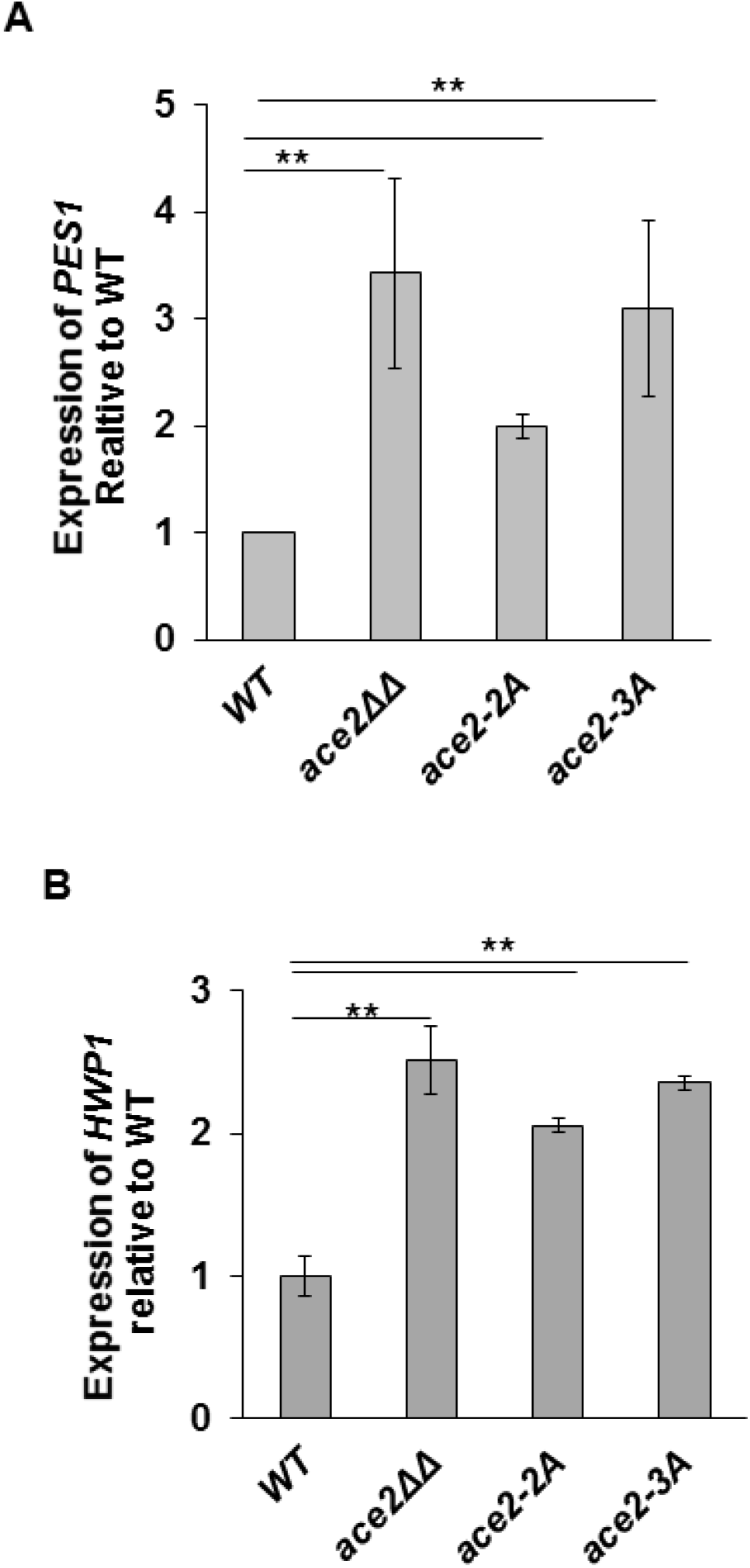
The expression of markers of *PES1* and *HWP1* is increased in *ace2*ΔΔ and *ace2-2A/3A* mutants. The indicated strains were induced to form hyphae in Spider medium (SM) at 37°C for 4 hours, harvested, and RNA isolated. The expression of *PES1* (A) and *HWP1* (B) were determined by quantitative RT-PCR and normalized to *ACT1* in each mutant. The relative expression of each gene was further normalized to WT. The bars indicate mean values for each strain with error bars indicating the SD of three independent experiments with technical replicates. The expression of each gene was statistically different than WT (Student’s t test, *P*<0.05).

## Discussion

The RAM pathway regulates a wide range of processes in *C. albicans* including cell cycle-associated daughter cell separation, hyphal morphogenesis, cell wall integrity and biosynthesis, biofilm formation, oxidative stress resistance, and mammalian infection (14–20). Although the phosphorylation motif of the key kinase in the RAM pathway, Cbk1, has been well-described (10), the substrates that carry out the effector functions of the pathway remain largely uncharacterized in *C. albicans* with the exceptions of Bcr1, Ssd1, and Fkh2 (27, 28, 29). For example, prior to the work described above, it was unclear what functions of the RAM pathway were due to its eponymous transcription factor Ace2.

Through transcriptional profiling of both *cbk1*ΔΔ and *ace2*ΔΔ mutants and genetic analysis of strains containing mutations at the Cbk1 phosphosites, we have found that, although both Cbk1 and Ace2 affect the expression of a large number of genes and have a number of phenotypes, the Cbk1-Ace2 axis directly regulates a specific subset of these functions. During yeast phase growth, the expression of only ~10% genes are concordantly affected in *cbk1*ΔΔ and *ace2*ΔΔ mutants. The cell septum degrading enzymes *CHT3* and *SCW11*, well characterized effectors of the RAM pathway (7), are among these and are also downregulated in Cbk1-phosphosite mutants of *ACE2*. As expected from studies of Cbk1 and Ace2 in *S. cerevisiae*, the *ACE2* phosphosite mutants have partial cell separation defects and alterations in the daughter cell specific nuclear localization localization of Ace2 (10).

The small overlap between the expression profiles of *cbk1*ΔΔ and *ace2*ΔΔ indicates that Cbk1 is likely to have additional transcriptional effectors and that Ace2 is likely to have other regulatory partners. With regard to the former, Bcr1 (27) and Fkh2 (28) have been shown to be Cbk1-regulated transcription factors. Bcr1 is important for biofilm formation but it’s effect on gene expression during yeast phase growth has not, to our knowledge, been characterized (36). Fkh2, interestingly, plays a role in both hyphal morphology and cell separation. During yeast phase growth, the expression of *SCW11* and *CHT2* is reduced in *fkh2* mutants (28). It is therefore possible that Cbk1 regulates cell separation genes through both Fkh2 and Ace2 and that other Cbk1-regulated genes are Fkh2 targets. A survey of the annotated *C. albicans* transcription factors indicates that an additional 27 have sequences that match Cbk1 phosphorylation motifs. Of these, a highly likely Cbk1 substrate is Ash1. Like Ace2 it is also a daughter cell-associated transcription factor that plays a role in hyphae formation under specific conditions. Its expression is also dependent upon both Cbk1 and Ace2 (Table S2&3). Ash1 has a HTRSR**S** Cbk1-consensus sequence in the N-terminus and S22 is phosphorylated during both yeast and hyphal phase growth according to two large-scale phosphoproteomic studies in *C. albicans* (30, 38).

It is also likely that another contributing factor to the role of Cbk1 in gene expression is its regulation of Ssd1, an RNA binding protein that suppresses the expression of a specific set of genes (29). In *S. cerevisiae*, *Sc*Cbk1 phosphorylates *Sc*Ssd1 to suppress its ability to target bound mRNAs to the P-body for storage and/or degradation (39, 40, 41). This set of *S. cerevisiae* genes is enriched in cell wall and cell separation genes but contains others as well (11, 39). In *C. albicans*, the only transcript that has been shown to be affected by Cbk1-Ssd1 is Nrg1 which is down-regulated at the initiation of hypha formation (29). It is unlikely that this is the only gene that is targeted by Ssd1, although additional studies will be needed to identify these transcripts. Overall, Cbk1 has a broad effect on *C. albicans* gene expression through a combination of its modulation of Ace2 activity, regulation of other transcription factors, and inhibition of Ssd1.

Ace2 also has broad effects on gene expression beyond cell separation as noted above (Fig. 1&2, Tables S1 and S3). Willger et al. found that Ace2 is phosphorylated at 16 sites in addition to those that match the Cbk1 motif, suggesting that other kinases are likely to be involved in the regulation of Ace2 function (30). Indeed, genetic studies by Van Wijlick et al. indicate that Ace2 functions in a kinase cascade comprised of Cst20, Hst7, and Cek1 to regulate protein mannosylation and susceptibility to the glycosylation inhibitor tunicamycin (21). Our data are consistent with their model in that the hypersusceptibility of *ace2*ΔΔ to tunicamycin is not recapitulated by *ACE2* mutants lacking Cbk1 phosphorylation sites (Fig. 6).

The Desai et al. have also performed ChIP-seq with Ace2 in YPD at 30°C under normoxia and hypoxia (17); because these data were not correlated with expression profiling, we compared the set of genes downregulated in the *ace2*ΔΔ mutant to the set of genes Desai et al. found to be bound by Ace2-HA under the same conditions. Surprisingly, only 23 of 359 (6.4%) of genes identified by Ace2-HA ChIP were down-regulated in the *ace2*ΔΔ mutant under these conditions. Further inspection of the ChIP dataset revealed that *SCW11*, *CHT3*, and *DSE3*, genes previously confirmed by single gene CHIP (42) were not among those identified as interacting with Ace2 during yeast phase growth. The reasons for the lack of overlap between these two large datasets are not clear.

Promoter motif analysis reported by Desai et al. identified a motif that was previously deduced by Mulhern et al. (16); however, only 11 of the genes reported by Mulhern et al. to be down-regulated in *ace2*ΔΔ were bound by Ace2-HA (17). One possibility is that the HA-tag is altering the function of Ace2 in a manner that is not evident from phenotypic analysis; although previous studies have shown that C-terminally tagged Ace2 proteins bind to targets such as *SCW11* and *CHT3* (42). It seems from these data and observations that additional experiments are needed before a definitive conclusion can be made regarding direct targets of Ace2 and the binding motifs that determine such targets.

Under many conditions, Ace2 is dispensable for *C. albicans* hyphal morphogenesis. Despite this, transcriptional profiling data reported by Mulhern et al. (16) and reported herein clearly indicate that it affects the expression of many genes during hyphal morphogenesis (Fig.2, Table S3). Our data indicate that a significant set of genes are upregulated in the absence of Ace2, suggesting that it functions either directly or indirectly as a repressor of gene expression during hyphal growth. Interestingly, *HWP1*, a gene that is only expressed during hyphae, and *PES1*, a gene associated with hyphae-to-yeast transition are both upregulated (Fig. 9A&B). *PES1* is essential during yeast phase growth and promotes the hyphae-to-yeast or lateral bud formation in hyphal cells (31); it is not essential for hyphal growth and forced expression drives lateral yeast cell formation in hyphae. Our data indicate that loss of Ace2 function and loss of Cbk1 regulation of Ace2 lead to increased expression of *PES1* and early formation of lateral yeast bud or hyphae-to-yeast transition (Fig.8&9). To our knowledge, this represents a novel function of Ace2 during morphogenesis and, as such, Ace2 is one of only a small set of genes that have been demonstrated to affect the hyphae-to-yeast transition (31, 39, 41).

We propose that the Cbk1-Ace2 axis functions during the maintenance phase of hyphae formation to suppress the lateral yeast growth program. During hyphal morphogenesis, Ace2 RNA and protein levels are initially low and then increase to peak at approximately 5 hours post induction in SM (32). Thus, except under specific conditions, Cbk1-regulated Ace2 does not appear to be required for initiation of hyphal formation. However, Ace2 appears to play an important role in sub-apical compartments by maintaining the hyphal transcriptional program or, alternatively, suppressing the yeast program (see below).

We and others have shown that Ace2-GFP is most clearly localized to the nuclei of the leading hyphal tip (15, 34). In addition to this well-established role in daughter cells, our data indicate that Cbk1-regulated Ace2 also functions in sub-apical hyphal cells as well. Although we have not been able to observe Ace2-GFP signal in the nuclei or cytoplasm of sub-apical hyphal cells (Wakade and Krysan, unpublished data), we suspect that this is due to the low overall expression of Ace2 from its native promoter; indeed, long exposure times are needed to visualize Ace2-GFP in the nuclei of daughter yeast cells or hyphal tip cells in our experience. The phenotypic data for the *ace2*ΔΔ and *ace2-2A/3A* strains, however, provide compelling evidence that Ace2 functions in the sub-apical and mother cell compartments of the hyphae.

Interestingly, our genetic data also indicate that the recently described Ace2L isoform may be required for this function (Fig. 3, 8 &9; reference 23). Because Calderón-Noreña DM et al. found that the Ace2S isoform is likely responsible for much of the transcriptional regulation attributed to Ace2 (23), it is possible that suppression of the yeast program is not a direct result of Ace2 binding to DNA; indeed, Ace2L is proposed to be membrane associated. Clearly, additional work will be needed to fully characterize the mechanistic details of this newly described Ace2 function.

Regardless of the specific molecular mechanism, the function of Ace2 in the sub-apical and mother cell compartments of the hyphal structure is consistent with its known cell cycle functions during yeast phase growth (9, 10) and the proposed cell cycle state of sub-apical compartments of *C. albicans* hyphae (6, 43). Specifically, Ace2 is well-established to play a critical role in early G1 during yeast phase growth in both *S. cerevisiae* (9, 10, 44) and *C. albicans* (45). As the cell progresses through G1 the amount of nuclear Ace2 is reduced by decreased *Sc*Cbk1 phosphorylation (44) and *Sc*Amn1-mediated ubiquitin-mediated degradation Ace2. This decrease in *Sc*Ace2 is associated with the transition to START (46). In contrast to yeast growth, Ace2 protein levels increase as hyphal morphogenesis progresses (32), suggesting that hyphal cells maintain an early G1-like state. Interestingly, *CaAMN1* expression is downregulated >3-fold relative to yeast cells after 4 hours of SM induction (Table S1 and S3), suggesting that decreased Amn1-mediated proteasome degradation of Ace2 could contribute to the steady increase in its protein level as hyphal morphogenesis progresses (32).

Sub-apical and mother cells of hyphae have been shown to be arrested in G1 phase (6, 44). Gow and colleagues have developed a model in which the highly vacuolated nature of the sub-apical compartments limits the amount of cytoplasm leading to an effectively smaller cell size (43, 48). Since cell size is a critical determinant of the cell transitioning from G1 to START, lateral yeast cell or branching does not occur until the ratio of cytoplasm to vacuole increases. Our data indicate that Cbk1-phosphorylated Ace2 plays a role in maintaining G1 and inhibiting the transition to START in both mother cells and sub-apical compartments. To our knowledge, this is the first genetic evidence for proteins that function to repress the transition from G1 to START in *C. albicans* hyphae.

Taking these new observations together with previous work from multiple labs including our own allows us to construct the following model integrating the function of the Cbk1-Ace2 axis during hyphal morphogenesis under standard liquid medium induction conditions (Fig. 10). After a hyphae-inducing signal is sensed, Cdc28 is activated (48) and, as demonstrated by Gutierrez-Escribano et al. (49), phosphorylates the critical activator of Cbk1, Mob2. Based on Lee et al., Cbk1 phosphorylates Ssd1 which, in turn, leads to degradation of the mRNA for the repressor of hyphal morphogenesis, Nrg1 (29); decreased Nrg1 activates hyphal morphogenesis through a number of pathways (50). Of the proteins de-repressed by decreased Nrg1 activity, the transcription factor Brg1 is particularly important (52). Indeed, Cleary et al. have shown that Nrg1 and Brg1 are part of a feedback loop that leads to increased *BRG1* expression during the initiation of hyphal morphogenesis (51).

**Figure 10.**
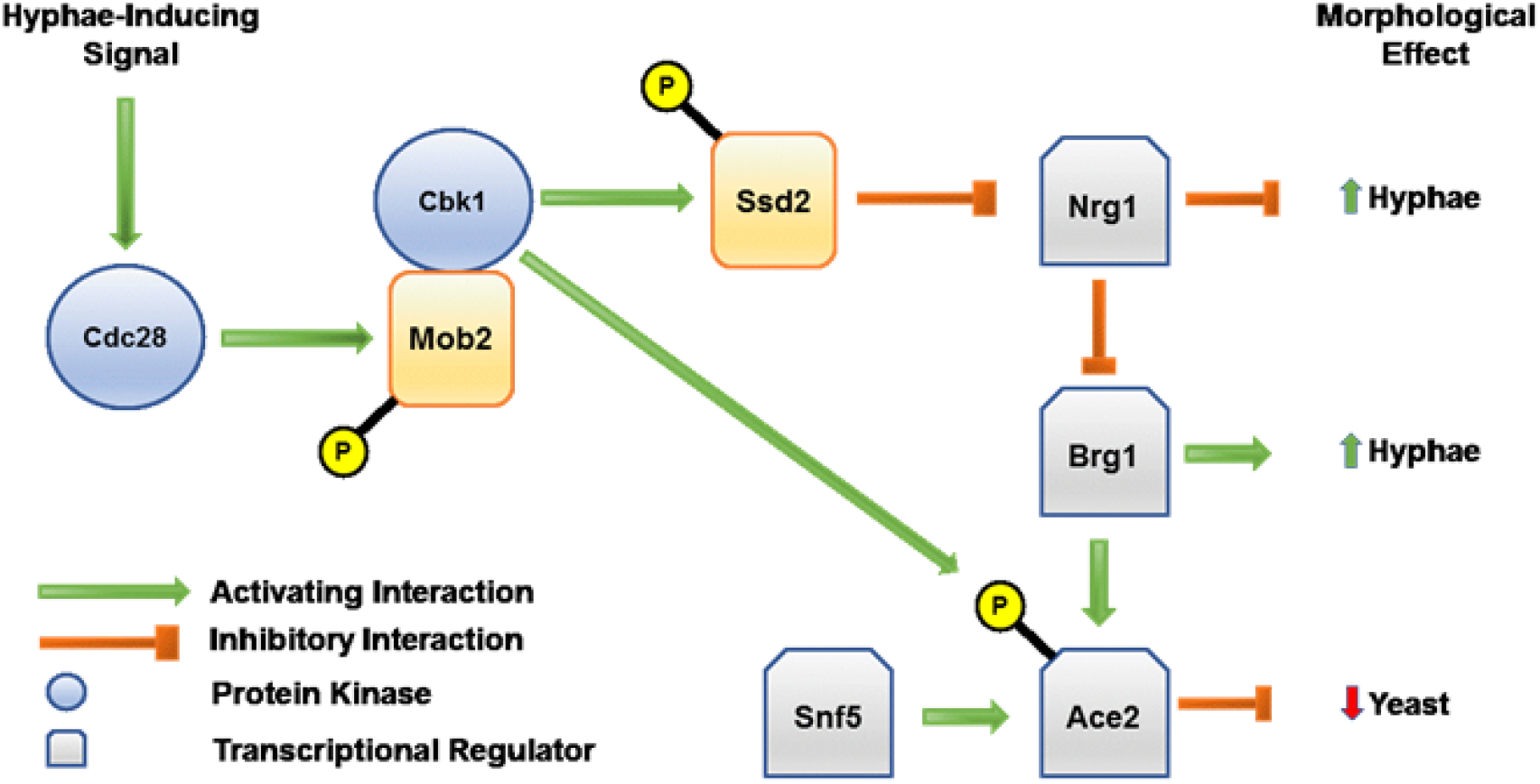
A model describing the function of the Cbk1-Ace2 axis during hyphal morphogenesis.

We have previously shown that Brg1 directly activates *ACE2* expression during hyphal morphogenesis and that expression peaks during the maintenance phase of the process (32). Finkel et al. found that Snf5 is also required for *ACE2* expression (18) and we show that it is also required for suppression of lateral yeast formation, indicating it plays a role in maintaining proper levels of *ACE2* during morphogenesis. Finally, as proposed above, Cbk1-phosphorylated Ace2 functions, at least in part, to suppress lateral yeast formation, likely through delaying the transition from G1 to START in subapical hyphal cell compartments.

In summary, Cbk1 regulates a specific set of Ace2 functions during both yeast and hyphae phase growth in *C. albicans*. While both Cbk1 and Ace2 have pleotropic effects on cell physiology, the majority of those effects are independent of one another. We have also proposed an integrated pathway through which Cbk1 affects hyphae formation during standard laboratory induction conditions which involves its regulation of the transcriptional regulators Nrg1, Brg1, and Ace2. Lastly, our data indicate that Cbk1-Ace2 is required to delay the G1-START transition in hyphae and, thereby, suppress the hyphae-to-yeast transition.

## MATERIALS AND METHODS

### Cultivation conditions, media, and materials

All *Candida albicans* strains were pre-cultured over-night in yeast peptone dextrose (YPD) medium at 30° C, unless indicated otherwise. Standard recipes (52) were used to prepare synthetic drop-out media, YPD, Spider medium (SM), and yeast peptone sucrose (YPS). Strains were induced for hyphal morphogenesis in liquid medium by diluting an over-night culture into SM and shifting to 37°C (32). Hyphal induction on solid surfaces was carried out either on SM (53, 54) or embedded in top agar-YPS medium following published protocols (16). Growth on YPD plates containing the indicated concentrations of calcofluor white, Congo red, and tunicamycin was examined as described (21). The chemical reagents were obtained from Millipore-Sigma and used as recieved.

### Strain construction

Strains and oligonucleotides used in this study are listed in the supplementary Tables S1 and S2, respectively. All heterozygous or homozygous strains of *C. albicans* were constructed from SN152 background using the auxotrophic marker either *LEU2* or *ARG4.* The *ace2Δ/ACE2* strain was generated as described previously (26). Briefly, pCR4-TOPO plasmid carrying *ACE2*::*LEU2* amplicon was digested with the SbfI enzyme and subsequently the linearized plasmid further inserted into the SN87 background (56). The transformants were selected on SD plates lacking leucine and correct integration was confirmed by PCR using ACE2.P1 and ACE2.P2 primers.

The transient CRISPR-Cas9 system was used to generate strains with deletion mutations (33). To generate *ace2ΔΔ* strains, a disruption cassette with 5’ and 3’ flanking sequences homologous to the 5’ and 3’ regions of *ACE2* was generated by amplification of *CmLEU2* cassette from pSN40 plasmid (56) with primer pairs ACE2.P3 and ACE2.P4. The *CaCas9* expression cassette was PCR amplified from plasmid pV1093 (33) and using CAS9.P1 and CAS9.P2 primers whereas sgRNA expression cassette was generated using the split-joint PCR method (33). Briefly, in the first step the SNR52 promoter was amplified using primer pairs ACE2.P5 and ACE2.P6 whereas sgRNA scaffold were PCR amplified by using ACE2.P7 and ACE2.P8 primers. In the second step, the SNR52 promoter and sgRNA scaffold were fused by primer extension in which the 20 bp guide RNA sequence act as complementary primers. In the third round, the fused PCR product was PCR amplified by using the nested primers ACE2.P9 and ACE2.P10 to harvest the sgRNA cassette. For fungal transformation, 1 μg of *CaCas9* cassette and 1 μg sgRNA cassette was co-transformed with the 3 μg of deletion construct, using the standard lithium acetate transformation method (33, 56).

To generate a strain with serine/threonine to alanine mutations at Cbk1 consensus phosphorylation sites, a fragment of *ACE2* (1-622 bp) that contained the respective mutations at T49A, S136A and S151A was synthesized by GeneScript and cloned into pUC19. This fragment was used as a template to generate strains containing the *ace2-2A* (S136A, S151A) and the *ace2-3A* (T49A, S136A, S151A) mutations. To do this, we used a double CRISPR approach. We first isolated *C. albicans ARG4* gene from a SC5314 background and PCR amplified using ARG4.P1 and ARG4.P2 primers. An *ARG4* cassette targeted to the putatively neutral *dpl200* locus was amplified using ARG4.P3 and ARG4.P4 primers. The phosphodeficient *ACE2* allele was amplified by using ACE2.P11 and ACE2.P12 primers. To achieve the transformation, 1 μg of *CaCas9* cassette, 1 μg of each sgRNA-ACE2 and sgRNA-*ARG4* cassette was co-transformed with the 3 μg of ACE2-2A/3A along with 1 μg of ARG4 cassette. The transformants were selected on synthetic media lacking Arg.

To characterize the resulting transformants, the region of *ACE2* corresponding to the repair fragment was PCR amplified using ACE2.P11 and ACE2.P12 primers and analyzed by Sanger sequencing. From multiple transformations, the majority of isolates were heterogeneous comprising of one copy of WT-ACE2 whereas other copy contained two (S136A, S151A) or three (T49A, S136A, S151A) mutations in *ACE2*. We then deleted the remaining WT copy of *ACE2* as described earlier (26) and the transformants were selected on SD plates lacking leucine and arginine. The correct integration was confirmed by PCR using ACE2.P1 and ACE2.P2 primers and the presence of the *ACE2* mutations was confirmed by Sanger sequencing. In this way, we generated strains in which the only copy of *ACE2* contained mutations in the Cbk1 phosphorylation sites.

Strains with GFP fused to the C-terminus of *ACE2* were generated by homologous recombination using a cassette that was derived from pMG2120 (57) and primers ACE2.P13 and ACE2.P14. The PCR amplified *NAT1*-marked, GFP cassette was purified and transformed into either the SN152 reference strain or *ACE2-2A* strain. Two independent clones of the respective strain were generated, and correct integration was confirmed by PCR. The resulting strains showed no change in growth or morphogenesis phenotypes relative to the parental strains. All PCR amplified or cloned fragments were confirmed by sequencing. Correct insertions of two independent clones were verified by PCR and used for the further experiments. Oligonucleotides were synthesized by IDT Technologies (Coralville, IA) and used as received.

### RNA sequencing

The indicated strains were grown overnight in YPD at 30° C and back diluted the next day and grown to exponential phase (WT, *ace2*ΔΔ and *cbk1*ΔΔ mutants) or induced to form hyphae with SM (WT and ace2ΔΔ strains) at 37°C for 4 hours. The cells were collected, centrifuged for 2 min at 11,000 RPM and RNA was extracted from the pellet according to the manufacturer protocol (MasterPure Yeast RNA Purification Kit, Cat. No. MPY03100). Briefly, the pellet was resuspended in 300 μL of Extraction reagent containing Proteinase K (50 μg/μl) and the mixture was incubated at 70° C for 15 min. The samples were placed on ice for 5 min and the cell debris was precipitated by addition of 175 μL of MPC reagent and centrifuged for 10 min at 11000 RPM at 4° C. The supernatant was transferred to the new tube and 500 μL of ice-cold isopropanol was used for mRNA precipitation. The obtained pellet was washed twice with ice-cold 70% ethanol, dried and mRNA was resuspended in 35 μL of TE buffer.

Total RNA submitted to the University of Wisconsin-Madison Biotechnology Center was verified for purity and integrity via the NanoDropOne Spectrophotometer and Agilent 2100 BioAnalyzer, respectively. Samples that met the Illumina sample input guidelines were prepared according the TruSeq^®^ Stranded mRNA Sample Preparation Guide (Rev. E) using the Illumina^®^ TruSeq^®^ Stranded mRNA Sample Preparation kit (Illumina Inc., San Diego, California, USA). For each library preparation, mRNA was purified from 1ug total RNA using poly-T oligo attached to magnetic beads. Following purification, the mRNA was fragmented using divalent cations under elevated temperature. The mRNA fragments were converted to double-stranded cDNA (ds cDNA) using SuperScript II (Invitrogen, Carlsbad, California, USA), RNaseH and DNA Pol I, primed by random primers. The ds cDNA was purified with AMPure XP beads (Agencourt, Beckman Coulter). The cDNA products were incubated with Klenow DNA Polymerase to add an ‘A’ base (Adenine) to the 3’ end of the blunt DNA fragments. DNA fragments were ligated to Illumina unique dual index Y-adapters, which have a single ‘T’ base (Thymine) overhang at their 3’ ends. The adapter-ligated DNA products were purified with AMPure XP beads. Adapter ligated DNA was amplified in a Linker Mediated PCR reaction (LM-PCR) for 10 cycles using Phusion™ DNA Polymerase and Illumina’s PE genomic DNA primer set followed by purification with AMPure XP beads. Finally, the quality and quantity of the finished libraries were assessed using an Agilent DNA1000 chip (Agilent Technologies, Inc., Santa Clara, CA, USA) and Qubit^®^ dsDNA HS Assay Kit (Invitrogen, Carlsbad, California, USA), respectively. Libraries were standardized to 2nM. Paired-end 2×150bp sequencing was performed, using standard SBS chemistry (v3) on an Illumina NovaSeq6000 sequencer. Images were analyzed using the standard Illumina Pipeline, version 1.8.2.

### Differential expression analysis

Paired-end Illumina sequence read files were evaluated for quality and the absence of adaptor sequence using FastQC (https://www.bioinformatics.babraham.ac.uk/projects/fastqc/). Read files were mapped to *C. albicans* reference genome SC5314 (FungiDB) and gene transcript expression was quantified using HISAT2 and Stringtie (58). Differential expression fold change, Wald test *P* values, and Benjamini-Hochberg adjustment for multiple comparisons were determined using DESeq2. Principle component analysis was performed on regularized, log transformed gene counts to confirm the absence of batch effects (59).

### Spot dilution phenotypic assays

The indicated strains were grown overnight in YPD at 30° C and back diluted the next day into the fresh YPD and grown for 4 hours at 30° C. The cells were further harvested and adjusted to cell density of 0.1 OD_600_ and tenfold serial dilution were performed and spotted on either YPD or YPD with the indicated concentration of calcofluor white, Congo Red, or tunicamycin. The plates were then incubated at 30° C and pictures were taken after 2 and 3 days of incubation. The assays were done in duplicate or triplicate on different days to confirm reproducibility.

### Assays of hyphal morphogenesis

For assays in liquid SM, strains were incubated overnight in YPD at 30°C, harvested, and diluted into SM at 1:100 ratio and incubated at 37° C for 4 hours. Cells were collected and either fixed with formaldehyde or examined by light microscopy directly. For assays on solid media, cells were grown overnight in YPD at 30° C, harvested and adjusted to cell density of 0.1 OD_600._ Ten-fold serial dilution was performed and cells from the fourth dilution were collected and plated on the indicated medium. For hypoxia phenotypes, the plates were incubated in chamber with the indicated amount of oxygen. For embedded experiments, cells were plated on to the YPS media and further overlaid with the YPS top agar and incubated at 25° or 37° C for 2 to 3 days.

### Microscopy

For colony morphology analysis, plates were incubated for the indicated time and imaged using Nikon ES80 epifluorescence microscope equipped with a CoolSnap charge coupled device (CCD) camera using 40x magnification. Live cell fluorescent imaging was carried out with the multi-photon laser scanning microscope (SP8, Leica Microsystems). Overnight cultures in YPD media were collected and back diluted either into the fresh YPD or in the hyphal inducing media to achieve either yeast or hyphal growth, respectively. Cells were then harvested, washed twice into 1x PBS and resuspended into 1x PBS prior to imaging. The samples were treated with Hoeschst 33342 and incubated for 15 minutes in the dark at room temperature followed by washing twice with PBS. For Hoechst 33342 dye excitation wavelength of 350 nm and emission wavelength of 461 nm was used whereas to visualize the GFP tag, excitation wavelength of 395 nm and emission wavelength of 509 nm was used, and sequential images were acquired using 25x water immersive objective lens with 3.32x zoom factor. Images were further analyzed by using the ImageJ software.

### Transcriptional analysis by quantitative reverse transcription-PCR (qPCR)

Strains pre-cultured overnight in YPD at 30°C, back-diluted into either fresh YPD or in hyphae-inducing media and collected after 4 to 5 hours of incubation either at 30° or 37° C. RNA was isolated using RiboPure ™ kit and reverse transcribed using an iScript cDNA synthesis kit (170-8891; Bio-Rad). The qPCR reaction was performed using IQ SyberGreen supermix (170-8882; Bio-Rad) and primers used in this study were listed in S3 table. Briefly, each reaction contained 10 μl of the SYBER Green PCR master mix, 0.10 μM of the respective primers and 150 ng of cDNA as a template in a total volume of 20 μl. Data analysis was performed using 2 ^−ΔΔCT^ method and *ACT1* was used as an internal control. Data reported here are the means from 3 independent biological replicates performed in a triplicate.

## ACKNOWLEDGEMENTS

This work was funded by NIH grants 1R01AI098450 (DJK) and 1R01AI33409 (DJK); the funders had no role in study design, data collection and interpretation, or the decision to submit the work for publication. We thank Manning Huang and Aaron Mitchell (Pittsburgh/University of Georgia) for helpful discussions regarding construction of the Ace2 mutants using CRISPR/Cas9 and for providing *snf5*ΔΔ mutants. We thank Scott Moye-Rowley (Iowa) and Melanie Wellington (Iowa) for helpful discussions.

## Author contributions

RW (designed and performed experiments; analyzed data; prepared figures; edited manuscript); LR (analyzed data; edited manuscript); MS (analyzed data); AK (designed experiments/analyzed data); DJK (experimental design; data analysis; prepared draft and edited manuscript).

## Supplemental Material

Supplementary Figure 1. The morphology of heterozygous *ace2*Δ/*ACE2* is not altered in comparison to the WT. (A) The indicated strains were spotted on the YPD plate and the morphology was assayed after 3 days of incubation at 30°C. (B) The *ace2*Δ /*ACE2* doesn’t play a role in cell separation defect. The indicated strains were quantified for cell separation defect. Shown here is the percentage of groups of cells vs the indicated strains. Bars are the mean ± SD of the three independent experiments. *n* = >150 cells. (C) The *ace2*Δ/*ACE2* is not hyper-resistant to CFW. Serial dilutions of indicated strains were spotted on YPD media (Ctrl) and YPD with CFW (50 μg/ml) and the images were taken after 2 days of incubation at 30°C. (D) The heterozygote *ACE2* mutant is not critical for hyphae formation under embedded conditions. The indicated strains were embedded in YPS agar at 25°C and images of colonies were taken after 2 days.

Supplementary Figure 2. The functionality of WT-Ace2-GFP and *ace2-2A-GFP* is not altered in comparison to the WT or untagged *ace2-2A* strain. (A) The indicated strains were spotted on YPD plates and the morphology was assayed after 3 days of incubation at 30°C. (B) The indicated strains were quantified for the cell separation defect. Shown here is the percentage of groups of cells vs the indicated strains. Bars are the mean ± SD of the three independent experiments. *n =* >150 cells.

Supplementary Table 1. Strains used in this work.

Supplementary Table 2. Oligonucleotides used in this work

Supplementary Table 3. RNA sequencing data for WT and *ace2*ΔΔ strains in YPD at 30°C.

Supplementary Table 4. RNA sequencing data for WT and *cbk1*ΔΔ strains in YPD at 30°C.

Supplementary Table 5. RNA sequencing data for WT and ace2ΔΔ strain in Spider Medium at 37°C.

Supplementary Table 6. List of genes for each Venn diagram category.

## REFERENCES

1. McCarty TP, Pappas PG. 2016. Invasive candidiasis. Infect Dis Clin North Am. 30:103–24.

2. Romo JA, Kumamoto CA. 2020. On commensalism of Candida. J Fungi 6:16.

3. Alves R, Barata-Antunes C, Casal M, Brown AJP, Van Dijck P, Paiva S. 2020. Adapting to survive: how *Candida* overcomes host-imposed constraints during human colonization. PLoS Pathog. 16:e1008478.

4. Desai JV. 2018. *Candida albicans* hyphae: from growth initiation to invasion. J Fungi 4:10.

5. Villa S, Hamideh M, Weinstock A, Qasim MN, Hazbun TR, Sellam A, Hernday AD, Thangamani S. Transcriptional control of hyphal morphogenesis in *Candida albicans*. FEMS Yeast Res 20:foaa005.

6. Sudbery PE. 2011. Growth of *Candida albicans* hyphae. Nat Rev Microbiol 9:737–748.

7. Saputo S, Chabrier-Rosello Y, Luca FC, Kumar A, Krysan DJ. 2012. The RAM pathway in pathogenic fungi. Eukaryot Cell 11: 708–717.

8. Nelson B, Kurischko C, Horecka J, Mody M, Nair P, Pratt L, Zougman A, McBroom LD, Hughes TR, Boone C, Luca FC. RAM: a conserved signaling network that regulates Ace2p transcriptional activity and polarized morphogenesis. Mol Biol Cell 14:3782–3803.

9. Sbia M, Parnell EJ, Yu Y, Olsen AE, Kretschmann KL, Voth WP, Stillman DJ. 2008. Regulation of the yeast Ace2 transcription factor during the cell cycle. J Biol Chem 283:11135–11145.

10. Mazanka E, Alexander J, Yeh BJ, Charoenpong P, Lowery DM, Yaffe M, Weiss EL. 2008. The NDR/LATS family kinase Cbk1 directly controls transcriptional asymmetry. PLoS Biol. 6:203.

11. Jansen JM, Wanless AG, Seidel CW, Weiss EL. 2009. Cbk1 regulation of the RNA-binding protein Ssd1 integrates cell fate with translational control. Curr Biol. 19:2114–2120.

12. Kurischko C, Kim HK, Kuravi VK, Pratzka J, Luca FC. 2011. The yeast Cbk1 kinase regulates mRNA localization via the mRNA-binding protein Ssd1. J Cell Biol 192:583–598.

13. Gógl G, Schneider KD, Yeh BJ, Alam N, Nguyen Ba AN, Moses AM, Hetényi C, Reményi A, Weiss EL. 2015. The structure of an NDR/LATS kinase-Mob complex reveals a novel kinase-coactivator system and substrate docking mechanism. PLoS Biol. 13:e1002146.

14. McNemar MD, Fonzi WA. 2002. Conserved serine/threonine kinase encoded by *CBK1* regulates expression of several hypha-associated transcripts and genes encoding cell wall proteins in *Candida albicans*. J Bacteriol. 184:2058–2061.

15. Kelly MT, MacCallum DM, Clancy SD, Odds FC, Brown AJ, Butler G. 2004. The *Candida albicans* Ca*ACE2* gene affects morphogenesis, adherence, and virulence. Mol. Microbiol. 53:969–983.

16. Mulhern SM, Logue ME, Butler G. 2006. *Candida albicans* transcription factor Ace2 regulates metabolism and is required for filamentation in hypoxic conditions. Eukaryot Cell 5:2001–2013.

17. Desai PR, van Wijlick L, Kurtz D, Juchimiuk M, Ernst JF. 2015. Hypoxic and temperature regulated morphogenesis in *Candida albicans*. PLoS Genet. 11:e1005447.

18. Finkel JS, Xu W, Huang D, Hill EM, Desai JV, Woolford CA, Nett JE, Taff H, Norice CT, Andes DR, Lanni F, Mitchell AP. 2012. Portrait of *Candida albicans* adherence regulators. PLoS Pathog. 8:e1002525.

19. Blankenship JR, Fanning S, Hamaker JJ, Mitchell AP. 2010. An extensive circuitry for cell wall regulation in *Candida albicans*. PLoS Pathog. 6:e1000752.

20. Song Y, Cheon SA, Lee KE, Lee SY, Lee BK, Oh DB, Kang HA, Kim JY 2008. Role of the RAM network in cell polarity and hyphal morphogenesis in *Candida albicans*. Mol. Biol. Cell 19:5456–5477.

21. van Wijlick L, Swidergall M, Brandt P, Ernst JF. 2016. *Candida albicans* responds to glycostructure damage by Ace2-mediated feedback regulation of Cek1 signaling. 102:827–849.

22. Ballou ER, Avelar GM, Childers DS, Mackie J, Bain JM, Wagener J, Kastora SL, Panea MD, Hardison SE, Walker LA, Erwig LP, Munro CA, Gow NA, Brown GD, MacCallum DM, Brown AJ. 2016. Lactate signalling regulates fungal β-glucan masking and immune evasion. Nat Microbiol. 2:16238.

23. Calderón-Noreña DM, González-Novo A, Orellana-Muñoz S, Gutiérrez-Escribano P, Arnáiz-Pita Y, Dueñas-Santero E, Suárez MB, Bougnoux ME, Del Rey F, Sherlock G, d’Enfert C, Correa-Bordes J, de Aldana CR. 2015. A single nucleotide polymorphism uncovers a novel function for the transcription factor Ace2 during Candida albicans hyphal development. PLoS Genet. 11:e1005152.

24. Saputo S, Norman KL, Murante T, Horton BN, Diaz Jde L, DiDone L, Colquhoun J, Schroeder JW, Simmons LA, Kumar A, Krysan DJ. 2016. Complex haploinsufficiency-based genetic analysis of the NDR/Lats kinase Cbk1 provides insight into its multiple functions in *Candida albicans*. Genetics. 203:1217–1233.

25. MacCallum DM, Findon H, Kenny CC, Butler G, Haynes K, Odds FC. 2006. Different consequences of *ACE2* and *SWI5* disruptions for virulence of pathogenic and non-pathogenic yeasts. Infect Immun. 74:5244–5248.

26. Glazier VE, Murante T, Koselny K, Murante D, Esqueda M, Wall GA, Wellington M, Hung CY, Kumar A, Krysan DJ. 2018. Systematic complex haploinsufficiency-based genetic analysis of *Candida albicans* transcription factors: tools and applications to virulence-associated phenotypes. G3 (Bethesda). 8:1299–1314.

27. Gutiérrez-Escribano P, Zeidler U, Suárez MB, Bachellier-Bassi S, Clemente-Blanco A, Bonhomme J, Vázquez de Aldana CR, d’Enfert C, Correa-Bordes J. 2012. The NDR/LATS kinase Cbk1 controls the activity of the transcriptional regulator Bcr1 during biofilm formation in *Candida albicans*. PLoS Pathog. 8:e1002683.

28. Greig JA, Sudbery IM, Richardson JP, Naglik JR, Wang Y, Sudbery PE. 2015. Cell cycle-independent phospho-regulation of Fkh2 during hyphal growth regulates *Candida albicans* pathogenesis. PLoS Pathog. 11:e1004630.

29. Lee HJ, Kim JM, Kang WK, Yang H, Kim JY. 2015. The NDR kinase Cbk1 downregulates the transcriptional repressor Nrg1 through the mRNA-binding protein Ssd1 in *Candida albicans*. Eukaryot Cell. 14:671–683.

30. Willger SD, Liu Z, Olarte RA, Adamo ME, Stajich JE, Myers LC, Kettenbach AN, Hogan DA. Analysis of the *Candida albicans* phosphoproteome. Eukaryot. Cell 14:474–485.

31. Shen J, Cowen LE, Griffin AM, Chan L, Köhler JR. 2008. The *Candida albicans* pescadillo homolog is required for normal hypha-to-yeast morphogenesis and yeast proliferation. 2008. Proc Natl Acad Sci USA 105:20918–20923.

32. Saputo S, Kumar A, Krysan DJ. 2014. Efg1 directly regulates *ACE2* expression to mediate cross talk between cAMP/PKA and RAM pathways during *Candida albicans* morphogenesis. Eukaryot Cell. 13:1169–1180.

33. Min K, Ichikawa Y, Woolford CA, Mitchell AP. 2016. *Candida albicans* gene deletion with a transient CRISPR/Cas9 system. mSphere 1:e00130–16.

34. Bharucha N, Chabrier-Rosello Y, Xu T, Johnson C, Sobczynski S, Song Q, Dobry CJ, Eckwahl MJ, Anderson CP, Benjamin AJ, Kumar A, Krysan DJ. 2011. A large-scale complex haploinsufficiency-based genetic interaction screen in *Candida albicans*: analysis of the RAM network during morphogenesis. PLoS Genet. 7:e1002058.

35. Lindsay AK, Deveau A, Piispanen AE, Hogan DA. 2012. Farnesol and cyclic AMP signaling effects on the hypha-to-yeast transition in *Candida albicans*. Eukaryot Cell 11:1219–1225.

36. Fanning S, Xu W, Solis N, Woolford CA, Filler SG, Mitchell AP. 2012. Divergent targets of *Candida albicans* biofilm regulator Bcr1 in vitro and in vivo. Eukaryot Cell 11:896–904.

37. Inglis DO, Johnson AD. 2002. Ash1 protein, an asymmetrically localized transcriptional regulator, controls filamentous growth and virulence of *Candida albicans*. Mol Cell Biol 22:8669–8680.

38. Beltrao P, Trinidad JC, Fiedler D, Roguev A, Lim WA, Shokat KM, Burlingame AL, Krogan NJ. 2009. Evolution of phosphoregulation: comparison of phosphorylation patterns across yeast species. PLoS Biol. 7:e1000134.

39. Hogan DJ, Riordan DP, Gerber AP, Herschlag D, Brown PO. 2008. Diverse RNA-binding proteins interact with functionally related sets of RNAs, suggesting an extensive regulatory system. PLoS Biol. 6:e255.

40. Kurischko C, Kim HK, Kuravi VK, Pratzka J, Luca FC. 2011. The yeast Cbk1 kinase regulates mRNA localization via the mRNA-binding protein Ssd1. J Cell Biol 192(4):583–598.

41. Veses V, Richards A, Gow NA. 2009. Vacuole inheritance regulates cell size and branching frequency of *Candida albicans* hyphae. Mol Microbiol 71:505–519.

42. Wang A, Raniga PP, Lane S, Lu Y, Liu H. 2009. Hyphal chain formation in *Candida albicans*: Cdc28-Hgc1 phosphorylation of Efg1 represses cell separation genes. Mol Biol Cell 29:4406–4416.

43. Barelle CJ, Bohula EA, Kron SJ, Wessels D, Soll DR, Schäfer A, Brown AJ, Gow NA. 2003. Asynchronous cell cycle and asymmetric vacuolar inheritance in true hyphae of *Candida albicans*. Eukaryot Cell 2:398–410.

44. Mazanka E, Weiss EL. 2010. Sequential counteracting kinases restrict an asymmetric gene expression to early G1. Mol Biol Cell 21:2809–2820.

45. Côte P, Hogues H, Whiteway M. Transcriptional program of the *Candida albicans* cell cycle. Mol Biol Cell 20:3363–3373.

46. Fang O, Hu X, Wang L, Jiang N, Yang J, Li B, Luo Z. (2018). Amn1 governs post-mitotic separation in *Saccharomyces cerevisiae*. PLoS Genet 14:e1007691.

47. Gow NA, Gooday GW. 1987. Cytological aspects of dimorphism in *Candida albicans*. Crit Rev Microbiol 15(1):73–78.

48. Wang Y. 2016. Hgc1-Cdc28-how much does a single protein kinase do in the regulation of hyphal development in *Candida albicans*? J Microbiol 54:170–177.

49. Gutiérrez-Escribano P, González-Novo A, Suárez MB, Li CR, Wang Y, de Aldana CR, Correa-Bordes J. 2011. CDK-dependent phosphorylation of Mob2 is essential for hyphal development in *Candida albicans*. Mol Biol Cell 22:2458–2469.

50. Lu Y, Su C, Liu H. 2014. *Candida albicans* hyphal initiation and elongation. Trends Microbiol 22:707–714.

51. Cleary IA, Lazzell AL, Monteagudo C, Thomas DP, Saville SP. 2012. *BRG1* and *NRG1* form a novel feedback circuit regulating *Candida albicans* hypha formation and virulence. Mol Microbiol 85:557–573.

52. Homann OR, Dea J, Noble SM, Johnson AD. A phenotypic profile of the *Candida albicans* regulatory network. PLoS Genet. 5:e1000783.

53. Wakade R, Labbaoui H, Stalder D, Arkowitz RA, Bassilana M. 2020. Overexpression of *YPT6* restores invasive filamentous growth and secretory vesicle clustering in a *Candida albicans arl1* mutant. Small GTPases 11:204–210.

54. Calera JA, Zhao XJ, Calderone R. 2000. Defective hyphal development and avirulence caused by a deletion of the *SSK1* response regulator gene in *Candida albicans*. Infect Immun 68:518–525.

55. Bassilana M, Arkowitz RA. 2006. Rac1 and Cdc42 have different roles in *Candida albicans* development. Eukaryot Cell 5:321–329.

56. Noble SM, Johnson AD. 2005. Strains and strategies for large-scale gene deletion studies of the diploid human fungal pathogen *Candida albicans*. Eukaryot Cell 4:298–309.

57. Gerami-Nejad M, Forche A, McClellan M, Berman J. 2012. Analysis of protein function in clinical *C. albicans* isolates. Yeast 29:303–309.

58. Pertea M, Kim D, Pertea GM, Leek JT, Salzberg SL. 2016. Transcript-level expression analysis of RNA-seq experiments with HISAT, StringTie and Ballgown, Nature Protocols 11:1650–1667.

59. Love MI, Anders S, Kim V, Huber W. 2015. RNA-Seq workflow: gene-level exploratory analysis and differential expression. F1000Research.doi:10.12688/f1000research.7035.

